# Retinoid X Receptor Signaling Mediates Cancer Cell Lipid Metabolism in the Leptomeninges

**DOI:** 10.1101/2024.10.13.618083

**Authors:** Xinran Tong, Jan Remsik, Jeanette Brook, Boryana Petrova, Lele Xu, Min Jun Li, Jenna Snyder, Kiana Chabot, Rachel Estrera, Isaiah Osei-Gyening, Ana Rita Nobre, Helen Wang, Ahmed M Osman, Alan Y.L. Wong, Mega Sidharta, Sofia Piedrafita-Ortiz, Branavan Manoranjan, Ting Zhou, Rajmohan Murali, Pierre-Jacques Hamard, Richard Koche, Ye He, Naama Kanarek, Adrienne Boire

## Abstract

Cancer cells metastatic to the leptomeninges encounter a metabolically-challenging extreme microenvironment. To understand adaptations to this space, we subjected leptomeningeal-metastatic (LeptoM) mouse breast and lung cancers isolated from either the leptomeninges or orthotopic primary sites to ATAC-and RNA-sequencing. When inhabiting the leptomeninges, the LeptoM cells demonstrated transcription downstream of retinoid-X-receptors (RXRs). We found evidence of local retinoic acid (RA) generation in both human leptomeningeal metastasis and mouse models in the form of elevated spinal fluid retinol and expression of RA-generating dehydrogenases within the leptomeningeal microenvironment. Stimulating LeptoM cells with RA induced expression of transcripts encoding *de novo* fatty acid synthesis pathway enzymes *in vitro*. *In vivo*, while deletion of *Stra6* did not alter cancer cell leptomeningeal growth, knockout of *Rxra/b/g* interrupted cancer cell lipid biosynthesis and arrested cancer growth. These observations illustrate a mechanism whereby metastatic cancer cells awake locally-generated developmental cues for metabolically reprograming, suggesting novel therapeutic approaches.

## Introduction

On entering a new environment, metastatic cancer cells encounter a variety of challenges, each imposing distinct constraints on the growth of these invaders^1,2^. Leptomeningeal metastasis (LM), or metastasis to the membranous coverings surrounding the central nervous system (CNS), provides an extreme example of this principle. The leptomeninges are comprised of the pia, arachnoid, and contain the circulating cerebrospinal fluid (CSF). The CSF is a notably hypoxic, low micronutrient, minimal amino acid, and lipid-scarce biological fluid^3,4^. Despite this anatomic isolation and nutritional scarcity, select cancer cells proliferate within this desolate microenvironment, leading to brisk accumulation of neurological deficits and death. Although any solid tumor may result in LM, in modern clinical practice, LM most commonly results from breast, lung, and melanoma primaries^5^.

Previous efforts from our lab uncovered cancer cell strategies to acquire limiting extracellular iron within the leptomeninges, including cancer cell co-option of the granulocyte LCN2/SLC22A17 high-affinity iron uptake system^6^. However, extracellular iron represents just one of a myriad of scarce nutrients within the leptomeningeal space. We hypothesized that it is unlikely that cancer cells would employ individual programs to overcome each of the limiting nutrients in the microenvironment singly; selection for such a large array of individual traits would be unlikely to occur on a time scale consistent with disease. Instead, it seems more probable that cancer cells might make use of preexisting programs to carry out the needed large-scale metabolic reprogramming.

Select non-transformed proliferating cells face constraints similar to metastatic cancer cells^7–9^. In the context of development, rapidly dividing cells must make up biomass prior to the advent of a closed circulation. In the mature vertebrate, metabolically nimble tissues, such as the liver, respond to extracellular cues to generate needed metabolic products^10^. We posited that cancer cells within the leptomeninges respond to local microenvironmental cues. These signals enable the metastatic cancer cell to access and repurpose preexisting developmental pathways to metabolically reprogram and overcome the nutrient scarce microenvironment of the leptomeninges. To address this hypothesis, we have examined epigenetic, transcriptional, and metabolic responses of leptomeningeal metastatic cancer cells from both mouse models and human tissues within a variety of microenvironmental contexts.

Here, we demonstrate that leptomeningeal metastatic cancer cells (LeptoM cells) are effectively metabolically reprogrammed by the leptomeningeal microenvironment.

Epigenetic and transcriptomic data together reveal evidence of Retinoid X Receptor (RXR) signaling in LeptoM cells when these cells inhabit the leptomeningeal space. We find evidence of local microenvironmental generation of retinoic acid (RA) in both mouse models and human disease in the form of elevated retinol levels in the CSF and increased leptomeningeal expression of RA-generating dehydrogenases in the setting of LM. Within LeptoM cells, we find that stimulating RXRs (encoded by *Rxra, Rxrb, Rxrg*) and Liver X Receptors (LXRs, encoded by *Nr1h2, Nr1h3*) induce transcription of genes encoding *de novo* fatty acid (FA) biosynthesis enzymes, including the central enzyme *Fasn*. Genetic knockout of *Fasn* in the LeptoM cells slowed cancer cell growth within the leptomeningeal space; triple knockout of *Rxra/b/g* completely arrested cancer cell growth in the space. In addition, LeptoM cells harboring *Nr1h2/3* or *Rxra/b/g* deletion both revealed substantial deregulation in lipid metabolism as characterized in both liquid and spatial mass spectrometry approaches, consistent with profound interruption of this arm of biosynthesis. Together, these observations reveal dynamic metabolic reprogramming in response to locally-generated developmental signals and suggest novel therapeutic targets to interrupt metastasis in the CNS.

## Results

### Cancer cells in the leptomeninges demonstrate evidence of RXR signaling

Leveraging the tools of iterative *in vivo* selection, we have previously generated cancer cells competent to enter into and survive within the leptomeningeal space, termed LeptoM cells^11,12^ (Figure S1A). To address how these selected cancer cells respond to microenvironmental cues, we implanted these cells into either the leptomeningeal space or into the orthotopic primary site: we injected mCherry-expressing LLC LeptoM and 4T1 LeptoM cells^12^ intracisternally or orthotopically to lung parenchyma for LLC or mammary fat pad for 4T1 LeptoM (Figure 1A). Faithfully recapitulating human disease, LeptoM cells exist in two phenotypic states *in vivo*: floating within the CSF, or adherent to the pial surface^13^. Reasoning that these two populations of cells encounter slightly different leptomeningeal microenvironments, we collected and analyzed these two groups of LeptoM cells separately. These freshly isolated LeptoM cells from different anatomical sites were subjected to Assay of Transposase-Accessible Chromatin sequencing (ATAC-seq) (Figure S1B). Initial Principal Component Analysis (PCA) demonstrated that both LLC and 4T1 LeptoM chromatin landscapes cluster based on their anatomical location (Figure S1C-D), suggesting a strong microenvironmental impact on LeptoM cell epigenetic state. Similarly, unsupervised *k*-means clustering using genome-wide differential peaks revealed distinct chromatin accessibility patterns in response to each microenvironment, in both LLC LeptoM (Figure 1B) and 4T1 LeptoM (Figure 1C) models. We next focused on epigenetic changes specific to the leptomeningeal space comparing open genomic region-associated genes in leptomeninges-residing cells to their orthotopic counterparts. Network enrichment analysis was notable for the dominance of development-related pathways in both models (Figure 1D). Interestingly, each model revealed subtly different enrichment patterns, likely related to their differing cellular origins (Figure S1E-F).

**Figure 1:**
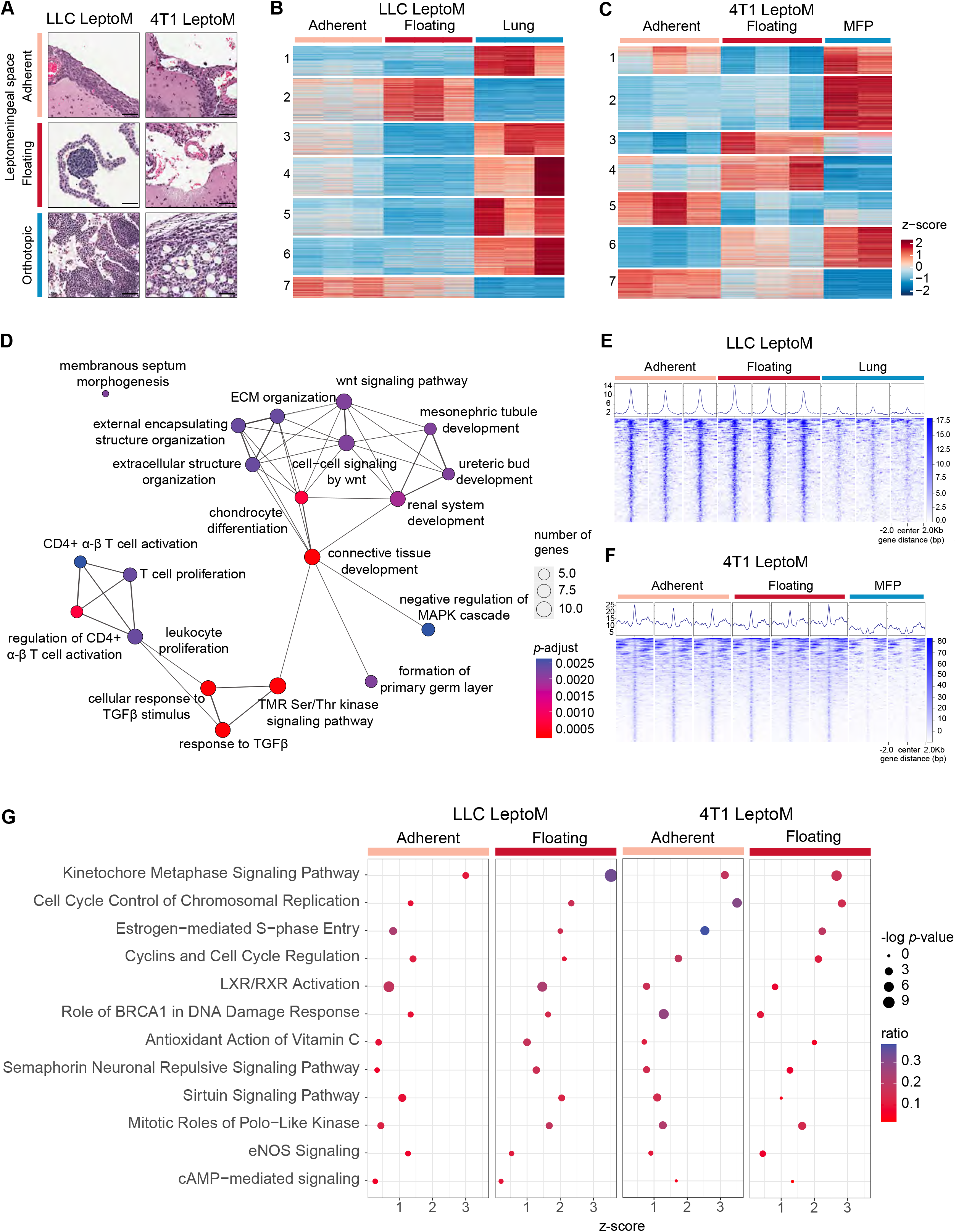
Epigenetic and transcriptomic analysis reveals evidence of RXR signaling in the leptomeninges. (A) H&E staining of tumors generated from LLC and 4T1 LeptoM cells instilled into different anatomic locations. Adherent: plaques of cancer cells adherent to the pial surface within the leptomeningeal space. Floating: Cancer cells captured floating within the ventricular system. Orthotopic: lung parenchyma for LLC LeptoM, mammary fat pad (MFP) for 4T1 LeptoM. Scale bars = 50μm. (B-C) Cancer cells collected from leptomeningeal space or the orthotopic primary site (please refer to Figure S1A-B) were subjected to bulk ATAC-seq. Genome-wide distribution of LLC and 4T1 LeptoM differential open chromatin peaks grouped by *k*-means clustering with bulk ATAC-seq signals. Each row represents one differential peak, normalized to sequencing depth, in sequential comparisons (log2|FC| > 2, FDR < 0.1). Each experimental group contains three technical replicates. 4T1 LeptoM harvested from MFP contains two technical replicates. (D) Genes associated with open chromatin in the leptomeninges vs. the orthotopic site in both LLC and 4T1 LeptoM models were subjected to combined network enrichment analysis. (E-F) Tornado plots of RXR motif-enriched genes in LLC LeptoM and 4T1 LeptoM. (G) Cancer cells collected from leptomeningeal space or the orthotopic primary site (please refer to Figure S1A-B) were subjected to bulk RNA-seq. (Please also refer to Figure S1H-K). Genes differentially expressed between the LeptoM adherent and floating fractions compared to their orthotopic counterparts populated the dataset and were subjected to ingenuity pathway analysis. Pathways in common between the LLC and 4T1 model systems with Z-score > 0.25 are depicted. LXR/RXR activation p-values are less than 0.027 across the groups.

We then focused on chromatin regions open in LeptoM cells inhabiting the leptomeningeal space, both floating and adherent, but absent in orthotopic sites consistently in both models, i.e., cluster 7 in *k*-means clustering heatmaps (Figure 1B-C). Motif binding prediction on cluster 7 revealed enrichment of the Retinoid X Receptor (RXR) motif in LLC LeptoM and Retinoid X Receptor/Retinoic Acid Receptor (RXR/RAR) motif in 4T1 LeptoM (Figure S1G). Indeed, we observed increased abundance of accessible RXR binding sites in cells inhabiting the leptomeningeal space in both models (Figure 1E-F). Taken together, the ATAC-seq analyses suggested a potential role for RXR in LeptoM cells residing in the leptomeningeal space.

To uncover the transcriptional consequences of this leptomeningeal-specific chromatin landscape change, we conducted high-throughput bulk RNA sequencing, again employing freshly isolated LLC and 4T1 LeptoM cells (Figure S1B). Consistent with the ATAC-seq results, the LeptoM transcriptome clustered based on microenvironment in both models (Figure S1H-I). Ingenuity pathway analysis (IPA) revealed concordantly upregulated pathways in floating and adherent populations in both models, including LXR/RXR activation (Figure 1G). Gene Set Enrichment Analysis (GSEA) uncovered enrichment of several metabolic pathways in leptomeningeal-residing cells related to RXR signaling. LLC LeptoM cells harvested from the leptomeninges displayed enrichment for steroid biosynthesis; 4T1 LeptoM displayed a signature of retinol metabolism (Figure S1J-K).

### LM provokes leptomeningeal generation of retinoic acid

Nuclear RXRs are activated by the short-lived molecule RA^14–16^, and function as transcription factors by forming homodimers or heterodimers with other nuclear receptors binding to different DNA direct repeats^17–19^. In the CNS, RXR signaling plays an essential role in nervous system development^20^. The ligand, RA is generated by the leptomeninges, and orchestrates cortical neuron generation^21,22^. We therefore pursued identification of RXR ligands within the leptomeningeal space, in the context of LM.

Originating from dietary sources, retinol is coupled to retinol binding proteins (RBPs), and circulates as retinol-RBPs^23^. RBPs mediate retinol delivery to target organ systems for local RA generation^24^. To synthesize RA, retinol is taken up by STRA6^25,26^, and oxidized to retinal primarily by retinol dehydrogenase 10 (RDH10)^27^. Subsequent irreversible conversion to RA is mediated by retinal dehydrogenase family proteins (RALDH1, RALDH2 and RALDH3), encoded by *ALDH1A1, ALDH1A2 and ALDH1A3* respectively (Figure 2A). The ephemeral, potent and freely diffusible RA is then available for local autocrine and paracrine activation of RXRs. To establish the scope of RA generation in the leptomeningeal space in the context of LM, we performed untargeted metabolomics using human cell-free CSF on Ultrahigh Performance Liquid Chromatography-Tandem Mass Spectroscopy (UPLC-MS/MS). Samples were selected from 35 newly diagnosed LM patients and 30 LM negative patients with breast, lung carcinoma, or melanoma primaries (Table S1). PCA based on 554 identified metabolites uncovered a trend of separation between LM negative and positive CSF samples (Figure S2A). Importantly, the dataset revealed a significant elevation in CSF *all-trans* retinol levels in the presence of LM (Figure 2B).

**Figure 2:**
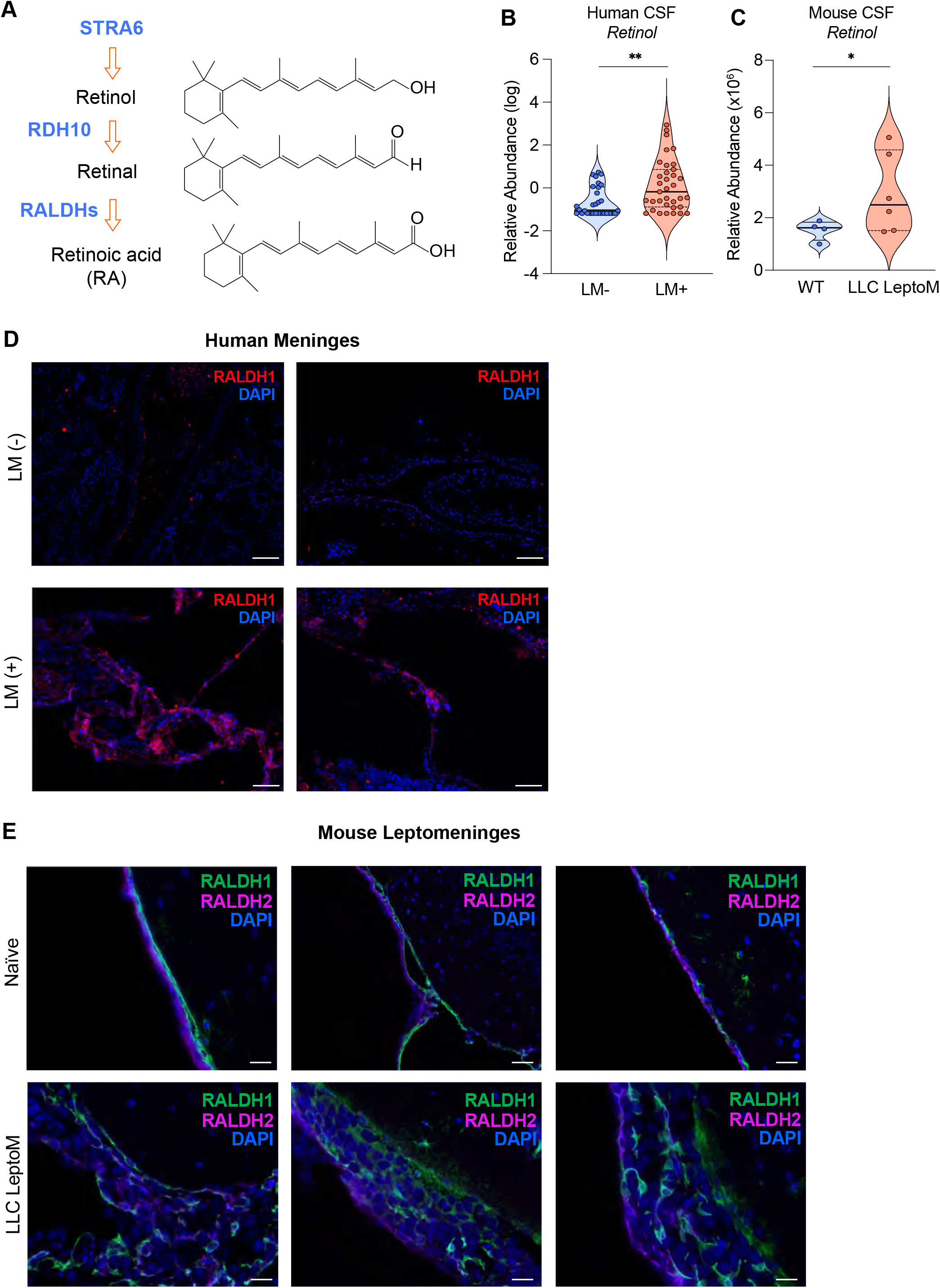
Increased retinol metabolism in the leptomeningeal microenvironment with LM. (A) Schematic drawing of human and mouse retinol metabolism pathway. Receptor and enzymes are blue, retinoids are black with chemical structure to the right. STRA6: Stimulated by Retinoic Acid 6; RDH10: Retinol Dehydrogenase 10; RALDHs: Retinaldehyde Dehydrogenases. (B) *All-trans* retinol level was detected in an untargeted UPLC-MS/MS in human cancer patient CSF without LM (n=30) or newly diagnosed with LM (n=35). Peak areas were batch normalized and imputed then log transformed and plotted. Black solid lines represent the median, dash lines represent quartiles. ** = 0.003, *p*-value represents *F*-test to compare population variance. (C) *All-trans* retinol level was detected on LC-MS/MS in the CSF from naïve mice (n=4) or mice injected with 2,000 LLC LeptoM intracisternally (n=6). Retinol standard was used for identification of retinol peak. Peak areas were normalized to a panel of polar metabolites (please refer to Figure S2B and Methods for details). Mouse CSF collected on day 14 after injection. * = 0.0458, *p*-value represents *F*-test to compare population variance. (D) Immunofluorescent staining of RALDH1 (red) and DAPI (blue) on human meningeal sections. Archival formalin-fixed paraffin-embedded sections of cancer patient meninges were obtained at autopsy with assigned identification code. Please also refer to Table S2. Two sections from patients without LM diagnosis or with LM diagnosis were selected. Scale bar = 100μm. (E) Immunofluorescent staining of RALDH1 (green), RALDH2 (magenta), and DAPI (blue) around the pia and arachnoid on mice brain sections collected from 3 naïve mice and 3 mice injected with 2,000 LLC LeptoM-mCherry cells intracisternally. LLC LeptoM cancer cells lie between pia mater and arachnoid layer. Scale bar = 20μm.

We next validated these findings in mouse models of LM. CSF was collected from naïve mice or mice harboring LM secondary to LLC LeptoM (Figure S2B). As in the human dataset, PCA based on 15 polar metabolites demonstrated moderate separation between naïve mice and mice harboring LM (Figure S2B). Consistent with human data, we observed elevated *all-trans* retinol in the CSF from mice with LM (Figure 2C); but not changed in mouse serum (Figure S2C). This is consistent with known impaired blood-CSF barrier integrity in the setting of LM and active retinol transport by the choroid plexus^11,28^.

Leptomeningeal fibroblasts orchestrate CNS development through local RA generation^21,29^. Indeed, prior work has established a meningeal RA source in the CNS in both embryonic and mature vertebrates^21,30^. Because malignant cells can provoke developmental and wound-healing responses in their local microenvironment^31–33^, we reasoned that these leptomeningeal cells may synthesize RA in the setting of LM. We therefore assayed RALDH1 expression by immunofluorescent staining in leptomeninges collected from cancer patients at autopsy (Table S2), comparing patients with and without LM (Figure 2D, S2D). In cancer patients without LM, the meningeal structures were intact, and RALDH1 was expressed sparsely throughout the meningeal tissues, predominantly in the arachnoid, consistent with prior observations^34^. In the presence of LM, the leptomeninges displayed structural disruption, as well as a higher level of RALDH1 expression throughout the leptomeninges.

In parallel, we observed similar findings in our mouse models. In mice, RALDHs are expressed in the meninges during development, and RALDH1 and RALDH2 expression is maintained into adulthood^35,36^. We found that leptomeninges from adult naïve mice demonstrated detectable levels of both RALDH1 and RALDH2, with RALDH1 expressed predominantly in the pia mater and RALDH2 in the arachnoid (Figure 2E). Expression of both proteins was preserved and elaborated in the leptomeninges of mice harboring LM. Consistent with local RA generation, LLC LeptoM cells adherent to the pia mater also expressed RALDH1 and RALDH2 proteins. To further identify evidence of retinol metabolism in human disease, we then interrogated our scRNA Seq dataset obtained from lung and breast cancer patients harboring LM^6^. This transcriptomic data demonstrated that cancer cells isolated from the spinal fluid transcribed a complement of genes involved in retinol uptake, mobilization, and metabolism (Figure S2E). This evidence of retinoid processing was phenocopied in our mouse models: in our previously published *in vitro* bulk RNA-seq datasets, LeptoM cell lines demonstrated increased expression of transcripts corresponding to retinol-binding protein 2 (*Rbp2*) in both LLC-LeptoM^11^ (Figure S2F) and 4T1-LeptoM^12^ (Figure S2G) models, as well as retinoic binding protein 2 (*Crabp2*) in 4T1 LeptoM, compared to their Parental (Par) counterparts.

Interestingly, in mice, expression of the dehydrogenase, Cytochrome P450 Family 1 Subfamily A Member 1 (*Cyp1a1*) was elevated in the setting of LM. This dehydrogenase can productively oxidize retinol direct to RA^37,38^. We observed elevated levels of CYP1A1 in choroid plexus epithelial cells of mice bearing LM compared to their naïve counterparts (Figure S2H). Taken together, these findings illustrate that CSF retinol can be processed into RA by relevant dehydrogenases by the leptomeningeal microenvironment, notably the arachnoid and pia to supply this ligand to the leptomeningeal space in the setting of LM.

### RXR stimulation induces *de novo* fatty acid biosynthesis in LeptoM cells

Previous work examining RXR heterodimerization activated by RA have highlighted the regulatory role of this signaling in lipid metabolism^39–41^. Among the nuclear receptors that partner with RXR, LXR/RXR activation regulates *de novo* fatty acid (FA) biosynthesis^42-^_44_. Both *all-trans* RA and *9-cis* RA serve as natural ligands for RXRs^14^, with *9-cis* RA exhibiting greater potency. To evaluate whether RXR stimulation induces upregulation of *de novo* FA biosynthesis in LeptoM cells, we treated LLC LeptoM cells with *all-trans* retinol, *all-trans* RA, or *9-cis* RA in delipidated fetal bovine serum supplemented media and assessed expression of genes encoding key enzymes responsible for *de novo* FA biosynthesis. Both 1μM *all-trans* RA and *9-cis* RA led to upregulation of *de novo* FA biosynthesis genes (*Srebf1, Acaca, Fasn, Scd1, Scd2*); treatment with *all-trans* retinol did not (Figure 3A). This pattern was also observed in B16 LeptoM cells (Figure S3A). Taken together, these results demonstrate that RA stimulation of RXR prompts expression of genes governing *de novo* FA biosynthesis in LeptoM cells. The upstream metabolite, retinol, cannot stimulate this transcription in these LeptoM cells alone, consistent with a key role for the microenvironment in this process.

**Figure 3:**
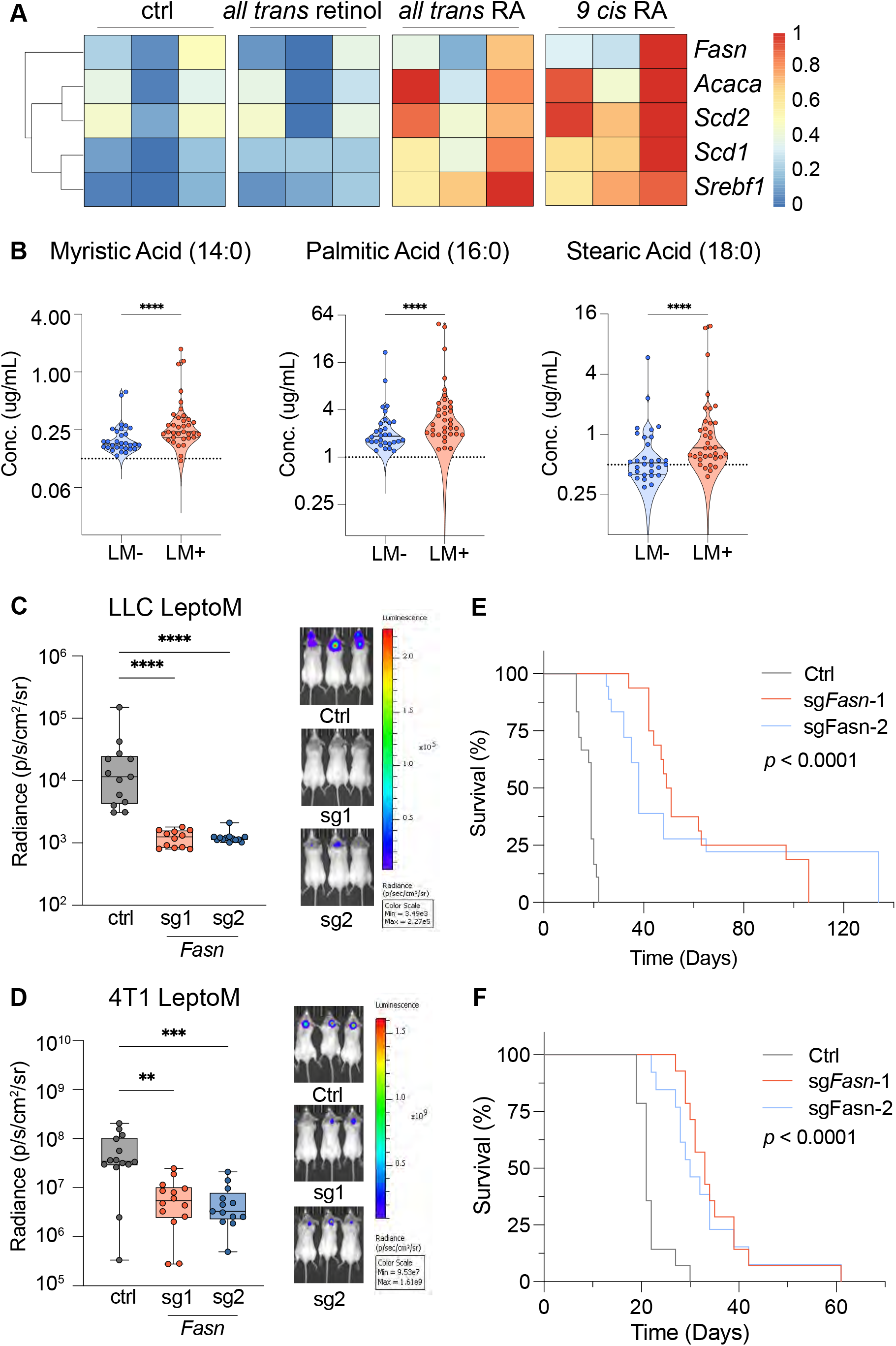
RA-activated *de novo* FA synthesis supports cancer cell growth in the leptomeningeal space. (A) RT-qPCR assaying *de novo* FA biosynthesis genes (*Srebf1, Fasn, Acaca, Scd1, Scd2)* in LLC LeptoM cells upon stimulation with 1μM *all-trans* retinol, 1μM *all-trans* RA, 1μM *9-cis* RA or vehicle control DMSO. Housekeeping genes Gapdh and *β*-actin were used as endogenous control. Stimulation in lipid-depleted media were normalized to LLC LeptoM DMSO-treated control in complete media. Plot contains three independent experiments each performed in duplicate. (B) Myristic acid (C14:0), palmitic acid (C16:0) and stearic acid (C18:0) were detected by targeted GC-MS lipidomics in human CSF samples (please refer to Figure 2B and Table S1). Concentrations were measured and plotted on log scale, dash lines represent lower limit of quantification. **** represents *p*-value <0.0001. *F-tests* were performed to compare population variance. (C) LLC LeptoM transduced with vector control or two independent small guide RNAs targeting *Fasn* were injected intracisternally into recipient mice. Tumor growth was monitored with bioluminescence imaging (BLI) weekly. The average radiance on day 14 was plotted, with representative images from each group. Ctrl: n=13; sg*Fasn*-1: n=12; sg*Fasn*2: n=13. Error bars represent SEM, *p*-value calculated by non-parametric one-way ANOVA (Kruskal-Wallis test), **** represents *p*<0.0001. (D) 4T1 LeptoM transduced with vector control or two independent small guide RNAs targeting *Fasn* were injected intracisternally into recipient mice; tumor growth was monitored by BLI weekly. Average radiance on day 14 is plotted, with representative images from each group. Three biological experiments were performed. Ctrl: n=14; sg*Fasn*-1: n=14; sg*Fasn*2: n=13. Error bars represent SEM, *p*-value calculated by non-parametric one-way ANOVA (Kruskal-Wallis test), ** represents *p*=0.0025, *** represents *p*=0.0007. (E) Kaplan-Meier survival curve of mice from (C). Ctrl: n=18; sg*Fasn*-1: n=16; sg*Fasn*2: n=18. Log-rank test was used for *p*-value. (F) Kaplan-Meier survival curve of mice from (D). Ctrl: n=14; sg*Fasn*-1: n=14; sg*Fasn*2: n=13. Log-rank test was used for *p*-value.

To further address the possible role of direct retinol stimulation of LeptoM cells, we investigated Stimulated by Retinoic Acid Gene 6 (STRA6), a membrane receptor and transporter for retinol^25^. We knocked out *Stra6* in 4T1 LeptoM with two independent small guide RNAs using CRISPR/Cas9 (Figure S3B). Intracisternal injection of 4T1 LeptoM *Stra6^-/-^* did not result in growth difference (Figure S3C) or consistent survival benefit (Figure S3D) compared to vector control. These results demonstrating lack of response to retinol alone illustrate the importance of leptomeningeal microenvironment to locally generate RA and enable transcription downstream of RXR.

Transcripts encoding multiple enzymes within the fatty acid biosynthesis pathway were downstream of RXR signaling in our system. To investigate the products of these enzymes, we conducted dedicated lipidomics to quantify free FAs in patient CSF samples (Table S1). We observed increased levels of both direct and indirect products of *de novo* FA biosynthesis in CSF collected from patients with LM, namely myristic acid (C14:0), palmitic acid (C16:0), stearic acid (C18:0) (Figure 3B). These data supported that RXR activation led to an upregulation of *de novo* FA synthesis in cancer cells *in vivo*.

### *De novo* fatty acid biosynthesis supports LeptoM cell growth in the leptomeninges

We hypothesized that cancer cells surviving in this low-lipid space are dependent on *de novo* FA biosynthesis. To address this, we knocked out expression of the gene *Fasn* encoding the central enzyme Fatty Acid Synthase by CRISPR/Cas9, with two independent small guide RNAs in LLC LeptoM (Figure S3E) and 4T1 LeptoM (Figure S3F). Leptomeningeal growth of LLC LeptoM (Figure 3C) and 4T1 LeptoM (Figure 3D) was significantly impaired in the *Fasn^-/-^* compared to their vector controls, resulting in prolonged survival in both models (Figure 3E-F). Another group observed similar cancer growth dependency on *de novo* FA synthesis in the breast cancer metastasis to the brain^4^.

We further assessed LLC and 4T1 LeptoM *Fasn^-/-^* cells within orthotopic primary sites. While FASN loss did impede LLC LeptoM cellular growth in the lung parenchyma (Figure S3G-H), it did not alter growth in 4T1 LeptoM within the mammary fat pad (Figure S3I). This inconsistency may well reflect lipid content differences between the lung parenchyma and mammary fat pad^45^. We reasoned that more robust interruption of this pathway might result from interference upstream of the individual enzymes of FA biosynthesis.

### LXRs and RXRs orchestrate *de novo* fatty acid biosynthesis in LeptoM cells

To confirm the upstream role of LXR and RXR in *de novo* FA biosynthesis *in vivo* and to assess functions of LXR and RXR in the LeptoM cells, we generated CRISPR/Cas9-mediated *Nr1h2* (encodes LXRα)/*Nr1h3* (encodes LXRα) double knockout (DKO) and *Rxra/b/g* (encode RXRα/α/ψ respectively) triple knockout (TKO) in LLC LeptoM cells. These genetic disruptions did not alter the cellular growth rate *in vitro* (Figure S4A). To address transcriptional implications of these alterations, we stimulated *Nr1h2/3* DKO, *Rxra/b/g* TKO, and vector control LLC LeptoM cells with 1μM *9-cis* RA or LXR synthetic agonist T0901317, then assessed transcription of *de novo* FA biosynthesis pathway genes by real time-quantitative polymerase chain reaction (RT-qPCR). In control cells, both agonists upregulated these genes (Figure 4A). *Nr1h2/3* DKO, demonstrated reduced upregulation of *de novo* FA biosynthesis pathway genes with T0901317 stimulation compared to the vector control. However, in this setting, the cells retained ability to respond to *9-cis* RA and transcribed *de novo* FA synthesis pathway genes, suggesting a compensatory mechanism through RXR, circumventing LXR. In contrast, *Rxra/b/g* TKO failed to upregulate *de novo* FA synthesis genes in response to either agonist, confirming that while LXR and RXR both function to regulate *de novo* FA biosynthesis, RXR serves a primary role for this function in this cellular context.

**Figure 4:**
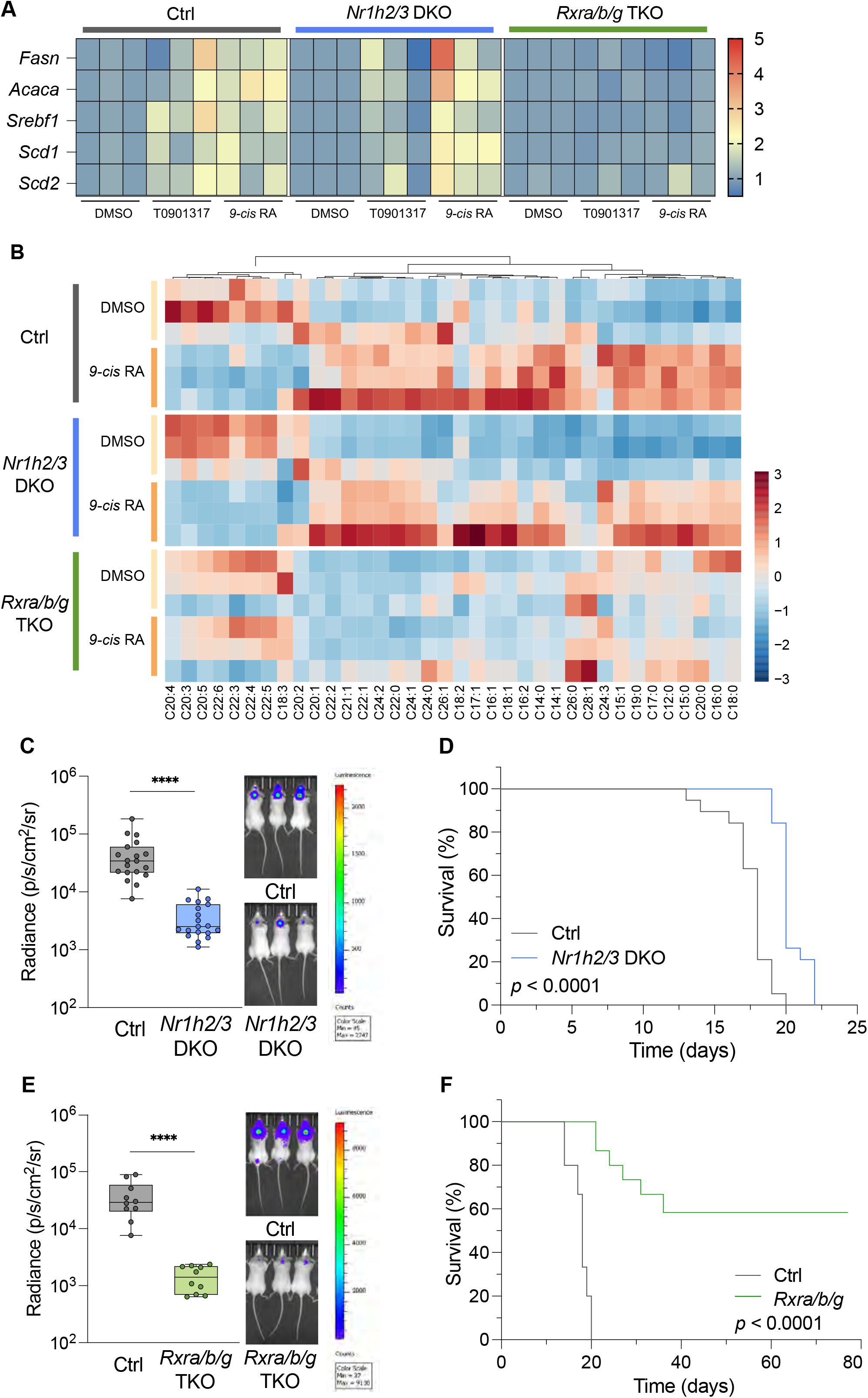
FA metabolism dysregulation in *Nr1h2/3* DKO and *Rxra/b/g* TKO. (A) RT-qPCR assaying *de novo* FA biosynthesis gene expression (*Srebf1, Fasn, Acaca, Scd1, Scd2)* upon stimulation with 1μM *9-cis* RA (RXR ligand), 1μM T0901317 (LXR ligand) or vehicle control DMSO in LLC LeptoM transduced with vector control, *Nr1h2/3* DKO or *Rxra/b/g* TKO. Housekeeping genes *Gapdh* and *β-actin* were endogenous controls. Experiments were performed in complete media with three independent experiments in duplicate. Each condition was normalized to its cell line DMSO control. (B) Intracellular free FA lipidomic analysis of LLC LeptoM vector control, *Nr1h2/3* DKO and *RXRα/α/ψ* TKO cells with DMSO vehicle control or 1μM *9-cis* RA. Peak intensities were normalized by median and scaled for heatmap. Three biological replicates are plotted as columns. (C) 2000 LLC LeptoM vector control or *Nr1h2/3* DKO cells were injected intracisternally into recipient mice; tumor growth was monitored by BLI weekly. The average radiance on day 14 was plotted, with representative images from each group. N=3 biological experiments; ctrl n=19; *Nr1h2/3* DKO n=19. Error bars represent SEM, *p*-value calculated by non-parametric T-test (Mann-Whitney test), **** represents *p*<0.0001. (D) Kaplan-Meier survival curve of mice from (C). *p*-value calculated by log-rank test. (E) 2000 LLC LeptoM vector control or *Rxra/b/g* TKO cells were injected intracisternally into recipient mice; tumor growth was monitored by BLI weekly. Average radiance on day 14 was plotted, with representative images from each group. N=2 biological experiments; ctrl n=15; *Rxra/b/g* TKO n=15. Error bars represent SEM, *p*-value calculated by non-parametric T-test (Mann-Whitney test), **** represents *p*<0.0001. (F) Kaplan-Meier survival curve of mice from (E). *p*-value calculated by log-rank test.

To correlate gene transcription with enzymatic function and further dissect the role of LXR and RXR in regulation of FA metabolism, both KO cell lines and vector control were stimulated with 1μM *9-cis* RA or DMSO control and subjected to lipidomic profiling for intracellular free FAs. In LLC LeptoM vector control cells, *9-cis* RA stimulation led to significant increase of intracellular free FAs, among which are palmitic acid (C16:0) and myristic acid (C14:0) (Figure 4B, Figure S4B). Additionally, odd-numbered carbon FA and long chain FA levels increased. Mammalian cells cannot synthesize these through *de novo* FA biosynthesis, indicating an upregulation in lipid uptake or lipid degradation with RA stimulation. In the LLC LeptoM *Nr1h2/3* DKO, RA stimulation led to exaggerated intracellular free FA changes (Figure S4C), with increased abundance of direct and indirect products from *de novo* FA biosynthesis. This confirmed our prior RT-qPCR observation whereby RA stimulation of *Nr1h2/3* DKO LeptoM cells led to an enhanced upregulation in *de novo* FA biosynthesis when compared to vector control (Figure 4A). This suggests a potential role for LXRs to monitor intracellular lipid levels and regulate FA biosynthesis in these cells. Changes in free FAs induced by *9-cis* RA were completely abolished in the LLC LeptoM *Rxra/b/g* TKO cells (Figure 4B), revealing the essentiality of RXR in upregulating *de novo* FA biosynthesis and potentially other lipid metabolic pathways.

### RXR-mediated signaling supports cancer cell growth in the leptomeninges

We next assessed the functional phenotype of these nuclear receptor KO cells *in vivo*. LLC LeptoM *Nr1h2/3* DKO grew less readily within the leptomeningeal space (Figure 4C) and resulted in an extension of survival (Figure 4D) compared to vector control group. In contrast, LLC LeptoM *Rxra/b/g* TKO, grew substantially slower in the leptomeningeal space (Figure 4E); a large proportion of the mice did not succumb to the disease (Figure 4F). Outside of the leptomeninges, loss of LXR (*Nr1h2/3* DKO) did not alter growth in subcutaneous (Figure S4D) or lung parenchyma (Figure S4E) sites, and did not alter survival (Figure S4F). However, *Rxra/b/g* TKO displayed growth disadvantage within both subcutaneous and lung parenchyma locations, as well as a substantial survival benefit for these mice (Figure S4D-F). Loss of function in RXRs halted the LLC LeptoM cellular growth to a much larger extent than *Fasn^-/-^* also matched with the complete ablation of FA changes upon RA stimulation in *Rxra/b/g* TKO. Taken together, these results underline the essential role of RXR in mediating *de novo* FA biosynthesis and supporting LeptoM cancer cell growth in the leptomeningeal space.

### The leptomeninges alters cancer cell fatty acid levels *in vivo*

The leptomeningeal space is remarkably compact in the rodent, approximately 36 microliters CSF in volume in an adult mouse^46,47^. *In vivo* assay of intracellular metabolites from these few LeptoM cancer cells is technically challenging. To capture LM-provoked FA metabolism changes *in vivo*, we performed matrix-assisted laser desorption/ionization mass spectrum imaging (MALDI MSI) on snap-frozen sections of LLC LeptoM injected mouse lung, LLC LeptoM vector control, or *Nr1h2/3* DKO injected mouse brain (Figure 5A). Hematoxylin and Eosin (H&E) staining highlighted the tumor cells located in the mouse lung parenchyma, or the leptomeningeal space surrounding the brain and ventricles. By mass spectrometry, glutathione [M-H]^-^ identified LLC LeptoM cancer cells in either the lung or leptomeningeal space; phosphatidylserine (PS) 36:1 [M-H]^-^ illustrated the brain structure. Further unbiased segmentation based on global spectra signatures distinguished tumor cells from adjacent normal tissues, confirmed by cross referencing H&E staining. We annotated and quantified four FAs, palmitic acid (C16:0), stearic acid (C18:0), oleic acid (C18:1), and arachidonic acid (C20:4), in the segmented tumor regions. All four FAs displayed higher mean intensities in the leptomeningeal-residing LLC LeptoM compared to the lung parenchyma-residing LLC LeptoM (Figure 5B, Figure S5A). In the LLC LeptoM *Nr1h2/3* DKO, mean intensities of the two *de novo* FA products, palmitic acid (C16:0) and stearic acid (C18:0), decreased compared to the LLC LeptoM control. In contrast, oleic acid (C18:1) and arachidonic acid (C20:4), derived from diverse intracellular sources, and not only FASN, were detected at comparable levels between LLC LeptoM *Nr1h2/3* DKO and LLC LeptoM control groups, in agreement with *in vitro* lipidomic data (Figure 4B). Curious to identify other leptomeningeal-specific metabolic changes, we carried additional unsupervised, high-resolution segmentation (Figure 5C, Figure S5B). We observed anatomic site-specific metabolic heterogeneity of the LLC LeptoM cells in both the lung and the leptomeningeal space (Figure 5D). This heterogeneity collapsed in the LLC LeptoM *Nr1h2/3* DKO population. Together, these data illustrate a distinct metabolic signature adopted by LLC LeptoM cancer cells and generation of *de novo* FA products in the leptomeningeal space.

**Figure 5:**
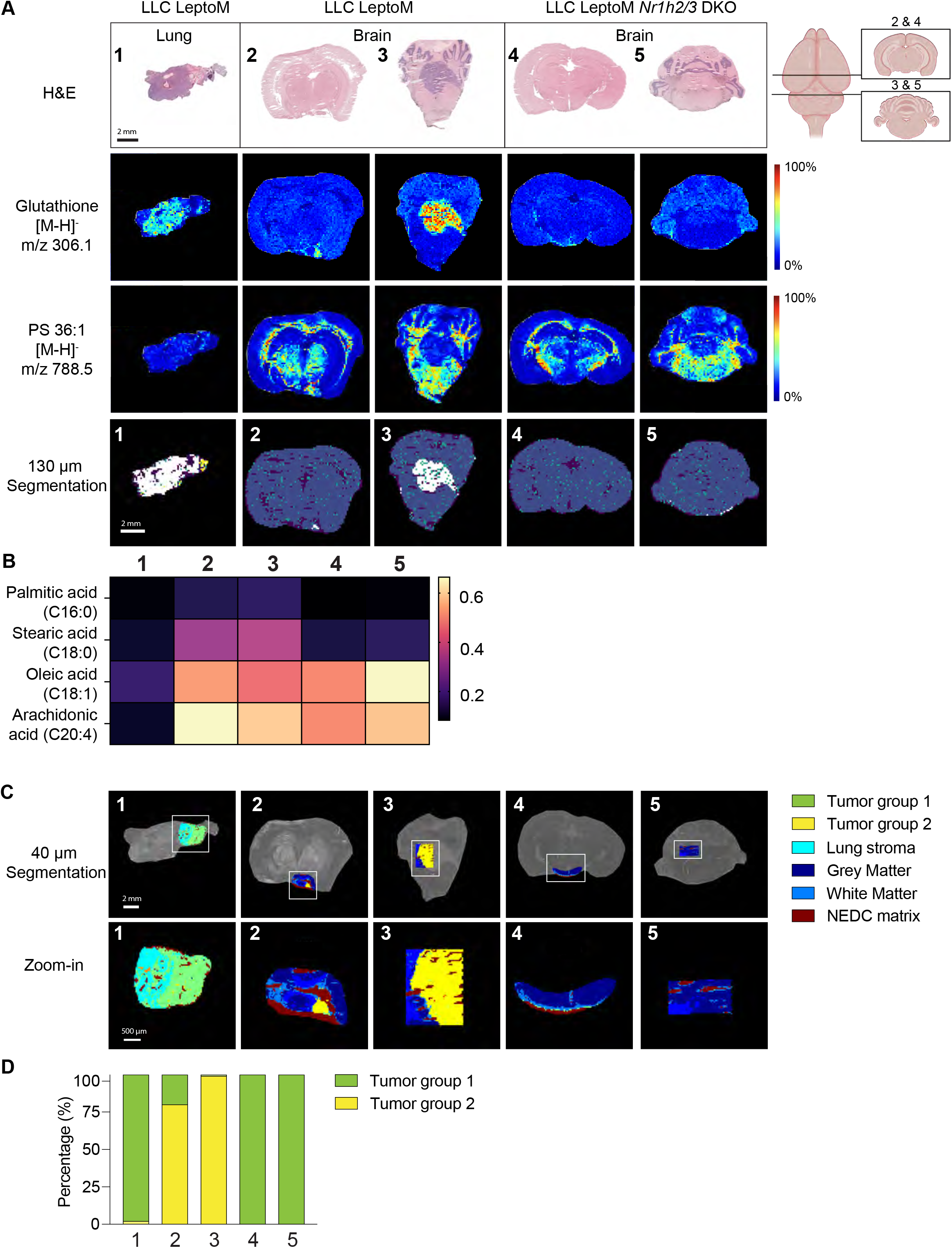
Spatial metabolomics confirm cancer cell leptomeningeal FA levels *in vivo*. (A) Untargeted matrix-assisted laser desorption/ionization mass spectrum imaging (MALDI MSI) on snap-frozen sections of LLC LeptoM injected mouse lung, LLC LeptoM vector control or *Nr1h2/3* DKO injected mouse brains. Each section is denoted with a number. Schematic drawings of mouse brain coronal sections cut at right. The tissue sections sprayed with NEDC matrix were subjected to MALDI MSI in negative ion mode. Sequential H&E staining together with reference metabolites (Glutathione [M-H]^-^ and Phosphatidylserine PS 36:1 [M-H]^-^) validated brain structures and cancer cells. Unbiased global spectra-based segmentation based on 130μm MALDI MSI highlights tumor regions in each tissue. Scale bar = 2mm. (B) Quantification of mean intensities for the identified fatty acids in the tumor region from 130μm segmentation. Each column corresponds to the numbered image in (A). (C) High-resolution 40μm scans on tumor and adjacent normal tissue sections (highlighted by white frames) with zoom-in images. Unbiased global spectra-based segmentation based on 40μm scans reveal two distinct subsets of tumor cells, colored as green (labelled as tumor group 1) and yellow (labelled as tumor group 2). Red region is identified as NEDC matrix background. Other colors are structures of the mouse brain. Details in Figure S5B and Methods. (D) Percentage of each tumor subset quantified in 40μm segmentation from (C) on each tissue section.

### Multifaceted role of RXR in lipid metabolism

Given the profound interruption of leptomeningeal cancer cell growth by RXR loss (Figure 4E-F), and the distinct metabolic signature adopted by cancer cells within the leptomeningeal space (Figure 5C-D), we hypothesized that RXR’s role in metabolic reprogramming of these LeptoM cells reached beyond *de novo* FA biosynthesis. Other than *de novo* FA biosynthesis, LXR and RXR are each known to play essential roles in lipid metabolism writ large (i.e. cholesterol biosynthesis, cholesterol efflux, and lipoprotein synthesis)^48–52^. Consistent with this, we observed a substantial elevation of structural lipids (phosphatidylcholine species and their associated lysolipid derivatives), as well as plasmalogens and sphingomyelin classes in CSF collected from cancer patients with LM compared to those without (Figure 6A, Table S3). To investigate LXR and RXR regulation on the larger cellular lipidome, we exposed the LLC LeptoM *Nr1h2/3* DKO, *Rxra/b/g* TKO and their vector control to *9-cis* RA or DMSO and subjected them to LC-MS/MS to assay intracellular complex lipid levels (Figure 6B). In vector control cell lines, RA stimulation led to deceased levels of cholesterol esters (CE) and sphingomyelins (SM) (Figure S6A). Significant enrichment of cardiolipins (CL) and phospholipids, namely phosphatidylglycerol (PG), phosphatidylinositol (PI), phosphatidylserine (PS), were observed in both nuclear receptor KOs compared to vector control (Figure S6B-C). *Nr1h2/3* DKO cells displayed decreased intracellular levels of diacylglycerols (DG) and hexosylceramides (HexCer) (Figure S6B), while *Rxra/b/g* TKO mainly exhibited a decrease in sphingomyelin (SM) with minor decrease in cholesterol esters (CE) and triglyceride (TG) (Figure S6C). As expected, RA stimulation in *Rxra/b/g* TKO did not elicit significant changes in intracellular complex lipids (Table S4). These results confirm that RXR exerts a master-regulator effect on lipid metabolism for LeptoM cells.

**Figure 6:**
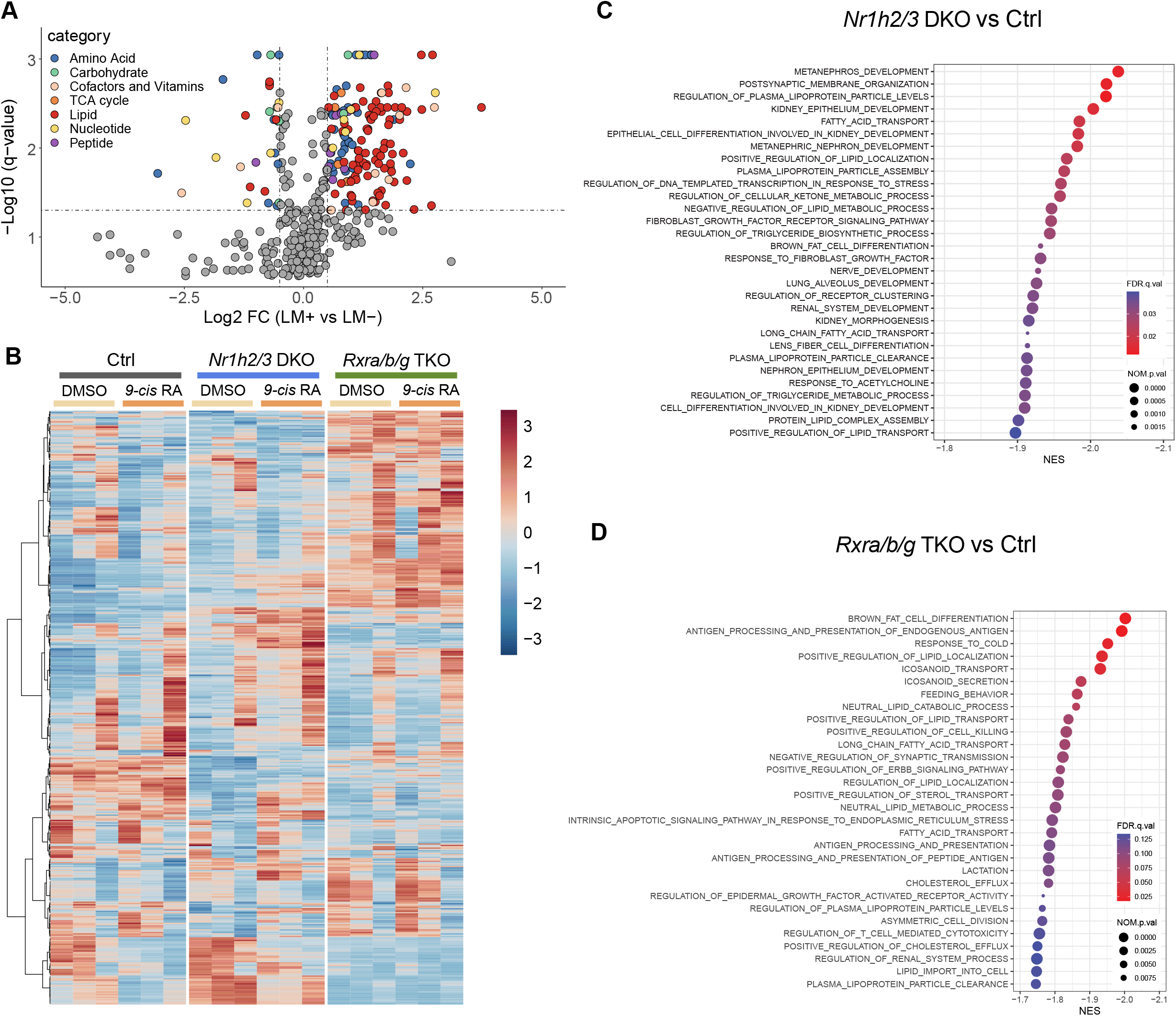
RXR regulates multiple aspects of lipid metabolism in LeptoM cells. (A) Volcano plot of untargeted metabolomics in human patient CSF comparing LM positive (n=35) to LM negative (n=30) (please refer to Table S1). Significantly changed metabolites are color-coded as categories. Significance cutoff: |log2 fold change| > 0.5, *q*-value < 0.05. (B) Intracellular complex lipid profiling heatmap performed by LC-MS/MS in LLC LeptoM vector control, *Nr1h2/3* DKO and *Rxra/b/g* TKO cells with DMSO vehicle control or 1μM *9-cis* RA *in vitro*. Peak intensities were normalized by median and scaled for heatmap. Three biological experiments were performed and plotted as columns. Each row represents a complex lipid detected. (C) IPA using significantly downregulated genes in LLC LeptoM *Nr1h2/3* DKO to vector control. Input differentially expressed gene dataset cutoff: *P.adjusted* <0.01, |log2FC| > 1. Top 30 significantly downregulated Gene Ontology_biological process (GO_BP) pathways were plotted. Pathway analysis cutoff: z-score <= -0.378, *p*-value <= 0.05. (D) IPA using significantly downregulated genes in LLC LeptoM *Rxra/b/g* TKO to vector control. Input differentially expressed gene dataset cutoff: *P.adjusted* <0.01, |log2FC| > 1. Top 30 significantly downregulated GO_BP pathways were plotted. Pathway analysis cutoff: *p*-value <= 0.05.

To gain insight into pathway-level alterations induced by these nuclear receptors, we subjected LLC LeptoM *Nr1h2/3* DKO, *Rxra/b/g* TKO and their vector control treated with *9-cis* RA or DMSO in complete media to bulk-RNA sequencing (Figure S6D). GSEA showed that RA stimulation in LLC LeptoM vector control triggered the activation of many developmental pathways (Table S5). Moreover, cholesterol biosynthesis (NES = 1.87, FDR = 0.046) (Figure S6E) and SREBF target (NES = 1.81, FDR = 0.082) (Table S5).

Pathways significantly enriched in the RA treated vector control cells align with our prior *ex vivo* ATAC-and RNA-sequencing signatures of LeptoM cell residing in the leptomeningeal space (Figure 1D, S1J-K). Many lipid metabolism pathways favoring complex lipid usage, such as neutral lipid metabolic process (NES = -2.08, FDR = 0.011) were significantly downregulated (Figure S6F, Table S5). As expected, RA stimulation in *Rxra/b/g* TKO led to significant changes in only 4 genes (Figure S6G), including *Dehydrogenase/Reductase 3* (*Dhrs3)*, which encodes an enzyme capable of reduction of retinal to retinol. We then focused on pathways downregulated after loss of LXR or RXR. GSEA revealed downregulation in several developmental pathways, lipid metabolism, transport and localization pathways in *Nr1h2/3* DKO (Figure 6C). In contrast, *Rxra/b/g* TKO exhibited downregulation in antigen presentation and immune-related pathways along with lipid metabolism pathways (Figure 6D). Commonly downregulated pathways shared between LLC LeptoM *Nr1h2/3* DKO and *Rxra/b/g* TKO cells were fatty acid transport, lipid localization, and lipoprotein particle levels. Of note, loss of RXR resulted in downregulation in cholesterol efflux pathways, consistent with decreased intracellular cholesterol esters detected in the *Rxra/b/g* TKO compared to control (Figure S6C). These findings collectively underscore the broad importance of RXR and LXR signaling in lipid metabolism beyond the narrow scope of FA metabolism in leptomeningeal metastatic cancer cells.

## Discussion

Metastatic cancer cells encounter novel microenvironments unlike those of their primary tumors, and must overcome these new local nutrient compositions to support their growth. In the leptomeninges, where there are multiple, significant sources of nutritional scarcity as well as a lack of capillaries for angiogenic support, these challenges take on major significance. Leptomeningeal metastasis therefore provides an ideal laboratory for the study of cancer cell metabolic reprogramming.

We have examined cancer cell epigenetic responses to the primary tumor microenvironment and the leptomeninges and uncovered evidence for metabolic plasticity. Plasticity, or the ability of cancer cells to change their phenotype in response to the environment, requires both a responsive epigenetic state in the cancer cell, and relevant signals from the microenvironment. Our analysis uncovered RXR signaling governing metabolic plasticity that is characteristic of LM. We demonstrated evidence of increased local leptomeningeal generation of RA in the setting of LM. We found that RA agonism of RXR with multiple heterodimer partners broadly upregulated lipid metabolism, while LXR/RXR activation promoted *de novo* FA biosynthesis. In support of this, we uncovered increased levels of lipids and products of *de novo* FA biosynthesis in LM patient CSF samples. Cancer cells within the space are reliant on lipid metabolism: loss of *Fasn* slows cancer cell growth; loss of RXRs arrests cancer cell growth. Compared to LXR, RXRs are known to have a broader impact on many aspects of cellular processes including development, differentiation, and metabolism^53^.

This dynamic metabolic reprogramming in response to extrinsic signals is reminiscent of the twin processes of development and wound healing. CSF contains morphogens and metabolites that are essential for brain development and CNS homeostasis^54^. RA represents a morphogen produced partially by the leptomeninges that governs CNS development^21,22^. It is intriguing to posit that RXR signaling might also result in metabolic reprogramming in the context of neural development. Individual isoforms of RXR are known to be essential to mediate development of certain tissues (i.e. RXRα in myocardial development and occulogenesis)^55,56^. Identification of the isoform(s) of RXR relevant for LM metabolic reprogramming will be helpful for generation of anti-LM therapeutics. However, RXRα can functionally substitute for the other two isoforms during development: the compound genetic knockout mouse, *RXRa^+/-^/RXRb^-/-^/RXRg^-/-^*, is viable^57^. While an array of agonists targeting RXR and select heterodimers is currently available, antagonists of the RXR remain less robustly developed^19^. Our genetic experiments support the key role that signaling downstream of this nuclear receptor plays in the pathogenesis of LM, and point to a novel therapeutic target.

In this work, we also provide an untargeted metabolomics database of human cancer patient CSF without LM or with newly diagnosed LM. This dataset encompassed three major solid malignancies that develop LM in modern clinical practice, and includes 645 species. This reference will serve as a valuable atlas for further study for a wide variety of investigators focused on metabolic vulnerabilities in metastatic cancer, as well as for those with interest in the composition and function of the blood-brain and blood-CSF barriers in health and disease.

### Limitations of the study

It remains unknown what purpose these FASN-derived lipids serve in the metastatic cancer cell, and is a major interest. The lipids might be employed directly for structural or signaling purposes-but might well be generated for an alternative purpose (i.e. to resolve a redox imbalance). Such work is beyond the scope of this current investigation.

## RESOURCE AVAILABILITY

### Lead contact

Further information and requests for resources and reagents should be directed to and will be fulfilled by the lead contact, Adrienne Boire.

## Materials availability

All cell lines generated in this study are available from the lead contact after completion of a Materials Transfer Agreement (MTA).

## Data and code availability

LLC and 4T1 LeptoM *ex vivo* bulk RNA- and ATAC-sequencing (GSE278626) and LLC LeptoM LXR and RXR KO *in vitro* bulk RNA-sequencing (GSE277622) have been deposited at GEO and are publicly available as of the date of publication. Accession numbers are listed in the key resources table. This paper also analyzes existing, publicly available data and can be retrieved from NCBI GEO with GSE150681). The accession numbers for the dataset are listed in the key resources table.

Mouse CSF targeted metabolite profiling analysis was supported by an in-house R script that performed technical and biological normalization and quality control steps and reported RSQ and coefficient of variation (CV). The code is shared here: https://github.com/FrozenGas/KanarekLabTraceFinderRScripts/blob/main/MS_data_scri pt_v2.4_20221018.R

Any additional information required to reanalyze the data reported in this paper is available from the lead contact upon request.

## EXPERIMENTAL MODEL AND STUDY PARTICIPANT DETAILS

### Animals

Animal studies were approved by the MSKCC Institutional Animal Care and Use Committee under protocol 18-01-002, and experiments were performed in accordance with this protocol. C57BL/6J (Stock #: 000664, Jackson Laboratory), B6(Cg)-Tyr^C-2J^/J (Stock #: 000058, Jackson Laboratory) and BALB/cJ (Stock #: 000651, Jackson Laboratory) were housed in the MSKCC vivarium, with individually ventilated cages, sterilized food and water. All mice were given at least one week to adjust to the habitat after transportation at above 5 weeks of age and used for experiments above 6 weeks of age. Murine lung cancer models were hosted in female and male mice at approximately 1:1 ratio. Murine breast cancer models were hosted in female mice. Mice of same sex were randomized before assignment to experimental groups. Mice used in the study were housed in controlled temperature and humidity, specific pathogen-free environment with 12-hour light/dark cycles (lights on/off at 6am/pm). Mice have access to standard chow and sterilized tap water *ad libitum*.

## Human Specimens

All human specimens were obtained with consent compliance with the MSKCC Institutional Review Board under MSKCC biospecimen protocols 20-117, 18-505, 13-039, 12-245, and 06-107. CSF samples were collected during routine neuro-oncologic care from consenting patients harboring breast, lung cancer or melanoma by lumbar punctures or through Ommaya reservoirs. 30 cancer patients without LM diagnosis at the time of collection were selected. 35 patients with LM were selected and the collections at diagnosis or the closest collections available after LM diagnosis were selected. Relevant clinical details are provided in Table S1. Freshly collected CSF samples were pelleted at 600 x g at 4°C for 5 mins in 1.5 mL Eppendorf tubes situated in 50mL conical tube in swing bucket centrifuge. Supernatant were carefully collected and aliquoted to store in - 80°C until sample analysis. Deidentified, archival formalin-fixed paraffin-embedded sections of cancer patient meninges were obtained from Last Wish Program with assigned identification code (Table S2). All above-mentioned patient gender, age, primary tumor diagnosis, leptomeningeal metastasis status, CSF cytology, CSF protein and glucose levels were recorded in Table S1 and S2.

## Cell Lines

Murine lung cancer derived LLC LeptoM, murine breast cancer derived 4T1 LeptoM and murine melanoma derived B16 LeptoM cell lines were described previously^11,12^. LLC and B16 LeptoM were maintained in high glucose DME medium (MSKCC media core), 4T1 LeptoM was maintained in RPMI 1640 medium (MSKCC media core). Both media were supplemented for a final concentration of 100U/mL penicillin-streptomycin (15140122, Gibco), 2mM GlutaMAX (35050061, Gibco) and 10% fetal bovine serum (FB-11, Omega Scientific). All cell lines were passaged at least twice a week and used under 20 passages, with routine mycoplasma tests. All cancer cell lines were engineered to express V5- tagged Firefly luciferase for tumor burden monitor and quantification. For bulk ATAC and RNA-seq, LLC LeptoM and 4T1 LeptoM were transduced with lentiviral particles to constitutively express mCherry (rLV.EF1.mCherry-9, Cat# 0037VCT, TakaraBio).

## METHOD DETAILS

### LeptoM Harvest for *ex vivo* Sequencing

Orthotopic and Intracisternal injections were previously described in detail^12^. LLC LeptoM mCherry were injected into mouse strain B6(Cg)-*Tyr^c-2J^*/J (The Jackson Lab) at 20,000 cells in 10uL PBS intracisternally or 25,000 cells in 50uL PBS into lung parenchyma. 4T1 LeptoM mCherry were injected into BALB/cJ (The Jackson Lab) at 5,000 cells in 10uL PBS intracisternally or 50,000 cells in 50uL PBS into the mammary fat pads. After two weeks, all mice were anesthetized with a mixture of 10mg/kg ketamine and 100mg/mL xylazine and perfused with PBS without Ca, Mg. Lung or mammary fat pad tumors were collected and dissociated with gentleMACS Dissociator (Miltenyi Biotec) using tumor program twice in PBS. To harvest leptomeningeal floating cells, surface of mouse brain, inner skull and lateral ventricle were rinsed with PBS. To harvest leptomeningeal adherent cells, rinsed mouse brains were submerged into 15mL falcon tubes containing an enzymatic cocktail (0.1% Dispase II, 0.01% Papain, 0.1% Collagenase type IV, 0.05% DNase I in 1x Hank’s balanced salt solution, no calcium, no magnesium supplemented with 12.4mM MgSO_4_) for 15mins at 37°C with gentle vertical rotation. All experimental groups were treated with the same enzyme cocktail for the same procedure to eliminate the bias on gene expression. Cells were washed, filtered and stained with a cocktail of FITC anti-mouse CD45 (1:200), FITC anti-mouse CD31 (1:200), FITC anti-mouse Ter- 119/erythroid cells (1:200), and 1ug/mL DAPI for fluorescence-activated cell sorting (FACS). For bulk RNA-seq analysis, mCherry-positive singlets were sorted into RLT buffer from RNeasy Plus Micro Kit with 1% α-mercaptoethanol added. All RNA were then isolated with RNeasy Micro Kit and further processed for high-throughput sequencing. For bulk ATAC-seq analysis, mCherry single positive cells were sorted into ice cold PBS and pelleted for downstream processing. About 3-5 mice were pooled for each technical replicate in all the experiments to get enough material for sequencing.

### ATAC-seq Processing and Analysis

Raw sequencing reads were trimmed for quality (Q≥15) and adapter content using version 0.4.5 of TrimGalore (https://www.bioinformatics.babraham.ac.uk/projects/trim_galore) and running version 1.15 of cutadapt and evaluated using FastQC (v0.11.5). Version 2.3.4.1 of bowtie2 (http://bowtie-bio.sourceforge.net/bowtie2/index.shtml) was used to align reads to mouse assembly mm10 with and duplicates were collapsed to one read using MarkDuplicates in version 2.16.0 of Picard Tools. Peaks were called using MACS2 (https://github.com/taoliu/MACS) with a p-value setting of 0.001, filtered for blacklisted regions (http://mitra.stanford.edu/kundaje/akundaje/release/blacklists/ mm10- mouse/mm10.blacklist.bed.gz), and a peak atlas was created using +/- 250 bp around peak summits. FeatureCounts (v1.6.1; http://subread.sourceforge.net) was used to build a raw counts matrix and DESeq2 was used to calculate differential enrichment (fold change ≥ 2 and FDR-adjusted p-value ≤ 0.1) for all pairwise contrasts. The sizeFactors from DESeq2 were used to normalize the raw counts matrix for clustering and heatmaps and for the creation of scaled bigwigs files using the BEDTools suite (http://bedtools.readthedocs.io). K-means clustering was performed on the superset of all differential peaks, increasing k until redundancy was observed. Peak-gene associations were created by assigning all intragenic peaks to that gene, while intergenic peaks were assigned using linear genomic distance to transcription start site. Gene Ontology and network analysis was performed on the genes assigned to differential peaks by running enrichGO and enrichplot::cnetplot in R with default parameters. Motif signatures were obtained using Homer v4.5 (http://homer.ucsd.edu) on differentially enriched peaks. Composite and tornado plots were created using deepTools v3.3.0 by running computeMatrix and plotHeatmap on normalized bigwigs with average signal sampled in 25 bp windows and flanking region defined by the surrounding 2 kb.

### Transcriptomic Analysis

All cancer cells collected *in vitro* were collected from 6-well plates with RLT buffer with 1% α-mercaptoethanol added and isolated by RNeasy Plus Mini Kit. All cancer cells from *in vivo* were collected as described above. mRNA isolated were used for library construction and sequenced by SMARTerAmpSeq with an average of 30-40 million reads per sample. The output FASTQ files were mapped to the *Mus musculus* genome (UCSC MM10) using the STAR aligner^58^ that mapped reads genomically and resolved reads across splice junctions. Two pass mapping method^59^ was used, then we computed the expression count matrix from the mapped reads using HTSeq (www- huber.embl.de/users/anders/HTSeq). The raw count matrix was then be processed for differential gene expression using the R/Bioconductor package DESeq2^60^. Pathway analyses were generated through the use of GSEA^61,62^ and QIAGEN IPA (QIAGEN Inc., https://digitalinsights.qiagen.com/IPA).

### Human CSF Metabolomics

#### Sample Preparation

Human CSF metabolomics analysis was performed at Metabolon Inc., Durham, North Carolina, USA. Samples were prepared using the automated MicroLab STAR® system from Hamilton Company. Several recovery standards were added prior to the first step in the extraction process for QC purposes. To remove protein, dissociate small molecules bound to protein or trapped in the precipitated protein matrix, and to recover chemically diverse metabolites, proteins were precipitated with methanol under vigorous shaking for 2 min (Glen Mills GenoGrinder 2000) followed by centrifugation. The resulting extract was divided into five fractions: two for analysis by two separate reverse phase (RP)/UPLC-MS/MS methods with positive ion mode electrospray ionization (ESI), one for analysis by RP/UPLC-MS/MS with negative ion mode ESI, one for analysis by HILIC/UPLC-MS/MS with negative ion mode ESI, and one sample was reserved for backup. Samples were placed briefly on a TurboVap® (Zymark) to remove the organic solvent. The sample extracts were stored overnight under nitrogen before preparation for analysis.

#### Ultrahigh Performance Liquid Chromatography-Tandem Mass Spectroscopy (UPLC-MS/MS)

All methods utilized a Waters ACQUITY ultra-performance liquid chromatography (UPLC) and a Thermo Scientific Q-Exactive high resolution/accurate mass spectrometer interfaced with a heated electrospray ionization (HESI-II) source and Orbitrap mass analyzer operated at 35,000 mass resolution. The sample extract was dried then reconstituted in solvents compatible to each of the four methods. Each reconstitution solvent contained a series of standards at fixed concentrations to ensure injection and chromatographic consistency. One aliquot was analyzed using acidic positive ion conditions, chromatographically optimized for more hydrophilic compounds. In this method, the extract was gradient eluted from a C18 column (Waters UPLC BEH C18- 2.1x100 mm, 1.7 µm) using water and methanol, containing 0.05% perfluoropentanoic acid (PFPA) and 0.1% formic acid (FA). Another aliquot was also analyzed using acidic positive ion conditions; however, it was chromatographically optimized for more hydrophobic compounds. In this method, the extract was gradient eluted from the same afore mentioned C18 column using methanol, acetonitrile, water, 0.05% PFPA and 0.01% FA and was operated at an overall higher organic content. Another aliquot was analyzed using basic negative ion optimized conditions using a separate dedicated C18 column. The basic extracts were gradient eluted from the column using methanol and water, however with 6.5mM Ammonium Bicarbonate at pH 8. The fourth aliquot was analyzed via negative ionization following elution from a HILIC column (Waters UPLC BEH Amide 2.1x150 mm, 1.7 µm) using a gradient consisting of water and acetonitrile with 10mM Ammonium Formate, pH 10.8. The MS analysis alternated between MS and data- dependent MS^n^ scans using dynamic exclusion. The scan range varied slighted between methods but covered 70-1000 m/z. Raw data files are archived and extracted as described below.

#### Data Extraction and Compound Identification

Raw data was extracted, peak-identified and QC processed using Metabolon’s hardware and software. These systems are built on a web-service platform utilizing Microsoft’s .NET technologies, which run on high-performance application servers and fiber-channel storage arrays in clusters to provide active failover and load-balancing. Compounds were identified by comparison to library entries of purified standards or recurrent unknown entities. Metabolon maintains a library based on authenticated standards that contains the retention time/index (RI), mass to charge ratio (*m/z)*, and chromatographic data (including MS/MS spectral data) on all molecules present in the library. Furthermore, biochemical identifications are based on three criteria: retention index within a narrow RI window of the proposed identification, accurate mass match to the library +/- 10 ppm, and the MS/MS forward and reverse scores between the experimental data and authentic standards. The MS/MS scores are based on a comparison of the ions present in the experimental spectrum to the ions present in the library spectrum. While there may be similarities between these molecules based on one of these factors, the use of all three data points can be utilized to distinguish and differentiate biochemicals. More than 3300 commercially available purified standard compounds have been acquired and registered into LIMS for analysis on all platforms for determination of their analytical characteristics. Additional mass spectral entries have been created for structurally unnamed biochemicals, which have been identified by virtue of their recurrent nature (both chromatographic and mass spectral). These compounds have the potential to be identified by future acquisition of a matching purified standard or by classical structural analysis.

#### Metabolite Quantification and Data Normalization

Peaks were quantified using area-under-the-curve. For studies spanning multiple days, a data normalization step was performed to correct variation resulting from instrument inter-day tuning differences. Essentially, each compound was corrected in run-day blocks by registering the medians to equal one (1.00) and normalizing each data point proportionately.

### Targeted Human CSF Lipidomics

Human CSF samples were analyzed by GC-MS at Metabolon, Inc. measuring the total content of fatty acids in human biological samples after conversion into their corresponding fatty acid methyl esters (FAME). For this assay margaric acid is also measured by conversion to the corresponding fatty acid methyl ester. This method was adapted for the analysis of cerebrospinal fluid and results are based on the analysis of 100 μL of human CSF sample. Quantitation ranges for the individual fatty acids are listed in Table 2. Aliquots of human CSF were pipetted into tubes and lyophilized. Internal standard solution was added to the lyophilized CSF samples. The solvent was removed by evaporation under a stream of nitrogen. The dried sample was subjected to methylation/transmethylation with methanol/sulfuric acid, resulting in the formation of the corresponding methyl esters (FAME) of free fatty acids and conjugated fatty acids. The reaction mixture was neutralized and extracted with hexanes. An aliquot of the hexanes layer was injected onto a 7890A/5975C GCMS system (Agilent Technologies, CA). Mass spectrometric analysis was performed in the single ion monitoring (SIM) positive mode with electron ionization. Quantitation is performed using both linear and quadratic regression analysis generated from fortified calibration standards prepared immediately prior to each run. Raw data are collected and processed using Agilent MassHunter GC/MS Acquisition B.07.04.2260 and Agilent MassHunter Workstation Software Quantitative Analysis for GC/MS B.09.00/ Build 9.0.647.0. Data reduction is performed using Microsoft Office 365 ProPlus Excel.

A single level QC was prepared for this project using pooled human cerebrospinal fluid. Pooled CSF was spiked with stock solutions to obtain the appropriate concentrations targeted at C to D level. Mean concentration values were set based on a ruggedness run prior to sample analysis. Sample analysis for the GC/MS was carried out in vials containing one calibration curve and six QC samples (per batch) to monitor assay performance. One batch of study samples was prepared and analyzed. QC accuracy was evaluated for the saturated fatty acids using the six QC replicates in the sample run. Intra-run accuracy for the saturated fatty acid analytes met acceptance criteria. (Targeted acceptance criteria for intra-run QC accuracy ≤ 20%.). Intra-run precision was also evaluated using the six QC replicates in the sample run. QCs met acceptance criteria for all saturated fatty acid analytes. Analyte concentrations that fell limit of quantitation are extrapolated and the cell is colored blue to indicate the sample is BLOQ.

### Mouse CSF and Blood Collection

Adult mouse CSF collection procedure is modified based on previous publications^63,64^. Mice were anesthetized with a mixture of 10mg/kg ketamine and 100mg/mL xylazine, then proned over a tip box to place the C-spine in flexion. The head area was prepared by removing the skin and cutting through the muscle along the midline to expose the cisterna magna. A pulled glass microcapillary tube was prepared by cutting a small opening. Glass microcapillary was introduced into the cisterna magna by avoiding blood vessels throughout the process. The microcapillary was connected to a 1TB syringe by a rubber tube. Once inside the leptomeningeal space, a second person pulled on the syringe extremely slowly to collect the CSF. Once CSF was in the capillary, the system was depressurized by pinching the tube and detaching the syringe. The capillary was carefully removed from the cisterna magna and the CSF was dispensed into protein low-binding tubes. Samples were then span down at 600g, 4°C for 5mins and supernatant was collected into ice-cold 80% methanol + 0.05% butylated hydroxytoluene (BHT) on dry ice and were immediately stored in -80°C for subsequent analysis. CSF from naïve group included 4 samples, and CSF from LLC LeptoM group included 6 samples with 2,000 LLC LeptoM cells injected intracisternally 2 weeks prior to collection. Blood samples were collected from left ventricles using 1TB 30G syringe with three naïve mice and three LM-baring mice. Serum was collected after blood clot at room temperature and collected into ice-cold 80% methanol + 0.05% BHT on dry ice and immediately stored in -80°C for subsequent analysis. All mice used were 8-week old male mice of the B6(Cg)-Tyr<C-2J= strain at the time of collection. All the procedure was done under yellow light to avoid degradation of retinoids. All tubes used were Protein Lo-Bind tubes. Methanol and H_2_O used were LC-MS grade.

### Mouse CSF Retinol Quantification by Mass Spectrometry

#### Extract retinoid metabolites and retinol detection

8-20 μL of CSF, plasma, and serum were combined with a ratio of 1:1:0.9 of chloroform:methanol:water supplemented with isotopically labeled internal standards (UltimateSPLASH™ ONE, retinol (R7632), metabolomics labelled amino acid mix). In addition, 0.05% BHT was included to minimize oxidation of retinoid metabolites during sample preparation. Samples were vortexed for 10 sec then centrifuged for 10 minutes at 18,000g to separate organic and aqueous phases. Each phase was transferred to a new 1.5 mL Protein LoBind Tubes and dried on ice in Reacti-Vap Evaporators using nitrogen flow. Metabolites were reconstituted in 15 μL of 75% methanol and 25% water supplemented with additional isotopically labeled internal standards, Metabolomics QReSS kit. For retinol detection, 8 μL of resuspended organic phase samples was injected into an Ascentis® Express C18 HPLC column (15 cm x 2.1 mm, particle size 2.7 μm; Sigma Aldrich) operated on a Vanquish™ Flex UHPLC System (Thermo Fisher Scientific, San Jose, CA, USA). The column oven and autosampler tray were held at 30 °C and 4 °C, respectively. The following conditions were used to achieve chromatographic separation: buffer A: water with 0.1% formic acid; buffer B: methanol. The chromatographic gradient was run at a flow rate of 0.250 ml min^−1^ as follows: 0–2 min: gradient was held at 75% B; 2–14 min: linear gradient of 75% to 98% B; 14.1–18.0 min: gradient was returned to 75% B. MS data acquisition was performed using a QExactive benchtop orbitrap mass spectrometer equipped with an Ion Max source and a HESI II probe (Thermo Fisher Scientific, San Jose, CA, USA). The mass spectrometer was operated in full-scan, positive ionization mode using one narrow-range scan: 260– 310 m/z, with the resolution set at 70,000, the AGC target at 1 × 10^6^, and the maximum injection time of 100 ms. HESI settings used were: sheath gas flow rate: 40 psi; aux gas flow rate: 10 a.u.; sweep gas: 0 a.u.; spray voltage: 3.5 kV (pos); capillary temperature 300 °C; S-lens RF level 50 a.u.; aux gas heater temp: 350 °C.

#### Polar Metabolite Detection for Normalization

1 μL of resuspended aqueous phase samples was injected into a SeQuant ZIC®-pHILIC column (150 x 2.1 mm, particle size 5 μm; Sigma Aldrich) operated on a Vanquish™ Flex UHPLC System (Thermo Fisher Scientific, San Jose, CA, USA). The column oven and autosampler tray were held at 25 °C and 4 °C, respectively. Chromatographic separation was achieved using the following conditions: buffer A: 95% acetonitrile, 5% HILIC mobile B (20 mM ammonium carbonate, 0.1% ammonium hydroxide in water; resulting pH is around 9.4); buffer B: 95% HILIC mobile B, 5% acetonitrile. The chromatographic gradient was run at a flow rate of 0.150 ml min−1 as follows: 0–20 min: linear gradient from 16.6% to 83.4% B; 20–24 min: hold at 83.4% B; 24.1-32 min: back to 16.6% B. MS data acquisition was performed using a QExactive benchtop orbitrap mass spectrometer equipped with an Ion Max source and a HESI II probe (Thermo Fisher Scientific, San Jose, CA, USA) in positive and negative ionization mode in a range of m/z = 70–1000, with the resolution set at 70,000, the AGC target at 1 × 10^6^, and the maximum injection time (Max IT) at 40 msec. HESI settings were: Sheath gas flow rate: 35 psi. Aux gas flow rate: 8 a.u. Sweep gas flow rate: 1 a.u. Spray voltage 3.5 kV (pos); 2.8 kV (neg). Capillary temperature: 320 °C. S-lens RF level: 50 a.u. Aux gas heater temp: 350 °C.

#### Metabolomics data analysis

Relative quantification of polar metabolites was performed with TraceFinder 5.2 (Thermo Fisher Scientific, Waltham, MA, USA) using a 10ppm mass tolerance and referencing an in-house library of chemical standards^65^. Pooled samples and fractional dilutions were prepared as quality controls and injected at the beginning and end of each run. In addition, pooled samples were interspersed throughout the run to control for technical drift in signal quality as well as to serve to assess the coefficient of variability (CV) for each metabolite. Data from TraceFinder was further consolidated and normalized with an in-house R script: (https://github.com/FrozenGas/KanarekLabTraceFinderRScripts/blob/main/MS_data_sc <u>ript_v2.4_20221018.R</u>). Briefly, this script performs normalization and quality control steps: 1) extracts and combines the peak areas from TraceFinder output .csvs; 2) calculates and normalizes to an averaged factor from all mean-centered chromatographic peak areas of isotopically labeled amino acids internal standards within each sample; 3) filters out low-quality metabolites based on user inputted cut-offs calculated from pool reinjections and pool dilutions; 4) calculates and normalizes for biological material based on the total integrated peak area values of high-confidence metabolites. In this study, the linear correlation between the dilution factor and the peak area cut-offs are set to RSQ>0.95 and the coefficient of variation (CV) < 30%. as well as to serve to access the coefficient of variability (CV) for each metabolite. Normalization of retinoid metabolites relied on retinol internal standard and a biological material normalizer based on polar metabolites. These steps were completed manually through Excel. First, retinoids peak areas were normalized to the mean-centered retinol internal standard (R-script step 2, technical normalizer). Second the resulting factor from the R script for polar metabolites (R-script step 4, biological normalizer), was used to account for any global shift in metabolite amounts due to differences in biological material.

### Single-cell analysis

The human single-cell dataset was retrieved from NCBI GEO as GSE150681^6^ and analyzed as described^12^.

### Human autopsy sample immunofluorescence

Human FFPE tissues were incubated at 60°C for 15mins, then deparaffinized and rehydrated with xylene (3 washes), 100% ethanol (2 washes), 95% ethanol (2 washes), 85% ethanol (2 washes), 70% ethanol and deionized water (2 washes) for 5 minutes each. Slides were incubated at 95°C in DAKO antigen retrieval buffer, pH =6, for 30 minutes, then cooled down slowly to room temperature. Permeabilization was done with PBS-T (0.1% Tween 20) + 0.4% Triton-X100 + 1% donkey serum for twice with 10 minutes each. Blocking was performed in PBS-T+5% donkey serum at room temperature for 1 hour. Primary antibodies were incubated in blocking buffer overnight at 4°C. Secondary antibodies were incubated in in blocking buffer at room temperature for 1 hour, then stained with 1ug/mL DAPI for 15 minutes. All washes were done in PBS-T. Slides were mounted with Prolong diamond antifade medium and scanned with Mirax slide scanner (Zeiss). Primary antibody: Goat anti-ALDH1A1, 1:40 dilution. Secondary antibody: Cy3 AffiniPure Donkey anti-Goat IgG (H+L), 1:800 dilution.

### Mouse immunofluorescence

Naïve mice or mice injected with LLC LeptoM intracisternally were anestisized and perfused with PBS, brains were harvested and fixed in 4% phosphate-buffered paraformaldehyde (FD Neuro Technologies, cat# PF101) for 48 hours at 4°C. Brains were then transferred to 30% sucrose at 4°C until they sank to the bottom. The brains were sliced into 4 cortical sections, placed in Tissue-trek vinyl cryomold 25x20x5mm (SAKURA, cat# 4557) with Tissue plus O.C.T. compound (Fisher HealthCare, cat# 4585) on dry ice. The tissues were then processed by MSKCC Molecular Cytology core facility, with cryosections at 7µm onto superfrost plus microscope slides and stored at -20°C until use. Cryosection slides were brought to room temperature and incubated at 60C for 30 mins. Slides were incubated in DAKO antigen retrieval buffer, pH =6, at 95°C for 30 minutes, then cooled down slowly to room temperature. Permeabilization was done with PBS-T (0.1% Tween 20) + 0.4% Triton-X100 + 1% donkey serum for twice with 10 minutes each. Blocking was performed in PBS-T+5% donkey serum at room temperature for 1 hour. Primary antibodies were incubated in blocking buffer overnight at 4°C. Secondary antibodies were incubated in in blocking buffer at room temperature for 1 hour, then stained with 1ug/mL DAPI for 15 minutes. All washes were done in PBS-T. Slides were mounted with Prolong diamond antifade medium and scanned with Mirax slide scanner (Zeiss). Primary antibody: goat anti-human/mouse ALDH1A1 (1:40), rabbit anti-human/mouse/rat ALDH1A2 (1:250), rabbit anti-human/mouse CYP1A1 (1:100), Goat anti-human/mouse/rat CD31/PECAM-1 (1:50). Secondary antibody: AF488 AffiniPure Donkey anti-rabbit IgG (H+L) (1:800), Cy3 AffiniPure Donkey anti-Goat IgG (H+L) (1:800).

### RT-qPCR for *de novo* fatty acid biosynthesis

Cells were plated at 0.2x10^5^ per well in 6-well plates on Day 0. Day 1, each well was rinsed with PBS, then switched to DMEM+10% FBS (Neuromics, FBS003) or DMEM+10% lipid-depleted FBS (Neuromics, FBS005-L). Day 2, 1uM *all-trans* retinol, *all-tra*ns RA, *9- cis* RA or same amount of Dimethyl sulfoxide control was added to the designated wells.

48hrs after treatments, RNA samples were collected and isolated with RNeasy Plus Mini Kit. 1500ng RNA from each sample was synthesized into cDNA with High-Capacity cDNA Reverse Transcription Kits, then assayed with Taqman Advanced Master Mix using Taqman probes: Srebf1, Fasn, Acaca, Scd1, Scd2. Housekeeping control: β-actin and Gapdh. Samples were run on QuantStudio 6 Flex machine (Applied biosystems), with three independent biological triplicates and technical duplicates each round. Data analyzed on QuantStudio Real-Time PCR software.

### LeptoM Cell Engraftment

Cells engineered under LLC LeptoM background were injected into B6(Cg)-*Tyr^c-2J^*/J at 2,000 cells in 10uL PBS intracisternally. For orthotopic injection, 25,000 cells in 50uL PBS were injected into lung parenchyma or 25,000 cells in 50uL PBS into subcutaneous location. Tumor burden was monitored with bioluminescence imaging (BLI) on IVIS Spectrum machine (Caliper Life Sciences) every 7 days, and morbidity was monitored and recorded daily. Quantification of tumor burden was analyzed by Living Image software (v4.7.3). Cells engineered under 4T1 LeptoM background were injected into BALB/cJ at 500 cells in 10uL PBS intracisternally or 50,000 cells in 50uL PBS into the fourth inguinal mammary fat pads. Mammary fat pad tumors were measured with digital Caliper (VWR, Cat# 62379531). Each experiment included at least two biological replicates, the exact numbers of round and mice were noted in the respective figure legends.

### Immunoblotting

Samples were lysed with RIPA Lysis and Extraction buffer with Halt Protease and Phosphatase Inhibitor Cocktail. Lysates were spun down at 13,000rpm in 4°C for 15mins, supernatant was collected into a new tube. Rapid Gold BCA protein assay kit was used to measure total protein levels in all samples. Samples were boiled in Laemmli buffer (Sigma Aldrich, #1610747) with 1% β-mercaptoethanol for 5 mins at 95°C. Proteins were resolved on a 4-15% Mini-PROTEAN TGC precast polyacrylamide gel (Bio-Rad, #4561083EDU) in 1x Tris/Glycine/SDS buffer (Bio-Rad, #1610772). Proteins were subsequently blotted onto 0.45μm PVDF membranes (Millipore Sigma, #IPVH00010) in 1x Tris/Glycine (Bio-Rad, #1610734) + 10% Methanol at 350A for 90 mins in 4°C, then blocked with 5% (w/v) fat-free milk powder (ChemCruz, #sc-2325) for 1hr at RT in Tris-buffered saline (Sigma Aldrich, #T5912-1L) with 0.01% Tween-100 (TBS-T). Subsequent incubation with primary antibody, human/mouse fatty acid synthase (R&D Systems, #MAB5927) at 1:500 dilution overnight at 4°C. β-Actin antibody (Millipore Sigma, # A5441-.2ML) at a dilution of 1:10,000 served as a loading control for total protein-level determinations. The secondary antibody employed was horseradish peroxidase-conjugated anti-mouse IgG (Cell Signaling, #7076S) at a dilution of 1:1,000. Immunoblots were developed with the ECL system (Pierce, #32109) and imaged with Amersham ImageQuant 800 (Cytiva).

### Generating genetic knockout of mouse LeptoM cancer cells

Plasmids with sgRNAs targeting mouse *Fasn* and *Stra6* and vector controls were generated by VectorBuilder. sgRNAs are expressed under the U6 promoter. *E.coli* stocks were expanded in LB with ampicillin overnight and plasmids were isolated with ZymoPURE II Plasmid kit (Zymo Research #D4203). Lentiviral particles were prepared with Lenti X 293T cell using VSV-G pseudotyped lentiviral packaging (TakaraBio, #631276) and concentrated with PEG gradient (TakaraBio, #631232). LeptoM cells were spin-transduced with concentrated lentivirus in the complete media with 10μg/mL polybrene (hexadimethrine bromide, Santa Cruz, #sc134220). After 24-48hr, 10μg/mL blasticidin (InvivoGen, #ant-bl-1) was added to allow for selection of edited clones. Once the cells recovered from selection, single-cell clones were generated either by single cell FACS or limited dilution into 96-well plates. Individual clones were screened either by western blots or genomic PCR following sanger sequencing.

### Generating LLC LeptoM *Nr1h2/3* DKO and *Rxra/b/g* TKO

sgRNA sequences were designed using Benching. Each target sequence was cloned into the lentiCRISPRv2 vector (Addgene, plasmid # 52961) to make lentivirus expressing sgRNA plasmids. The mouse cancer line LLC LeptoM cell line was infected with lentivirus carries the sgRNAs. For *Nr1h2/3* DKO, the cells were infected with *Nr1h2* and *Nr1h3* expressing sgRNA lenti-virus. For *Rxra/b/g* TKO the cells were infected with *Rxra*, *Rxrb* and *Rxrg* expressing sgRNA lenti-virus. Two days after infection, cells were treated with 1 μg/ml puromycin for 2 days. The cells were then dissociated into single cells and seeded as one cell one well into 96-well plates. Around 14 days later, individual colonies derived from single cells were picked, mechanically disaggregated, and replated into two individual wells of 96-well plates. A portion of the cells was analyzed by PCR and Sanger Sequencing. Biallelic frameshift mutants were chosen as knockout clones.

### LLC LeptoM *Nr1h2/3* DKO and *Rxra/b/g* TKO bulk RNA-sequencing *in vitro*

LLC LeptoM *Nr1h2/3* DKO, *Rxra/b/g* TKO and their vector control cells were plated with complete DME media at 0.2x10^5^ per well in 6-well plates on Day 0. After 24 hours, 1μM 9 cis RA or same percentage of DMSO were added to designated plates for 48 hours. Cells were lysed with RLT buffer from RNeasy Plus Mini Kit with 1% α-mercaptoethanol added. All RNA were then isolated with RNeasy Mini Kit and further processed for high-throughput sequencing. Two to three biological samples were included for each condition. Downstream processing was described in the transcriptomic analysis section.

### Lipidomics from *in vitro* LeptoM cells

#### Sample Preparation

1x10^6^ cells were seeded in 10cm plates. After 24 hours, 1μM 9 cis RA or same percentage of DMSO were added to designated plates for another 24 hours. At the time of harvest, each plate was trypsinized and counted with trypan blue on a Cellometer (Nexcelom Biosciences LLC, MA USA). 1x10^6^ cells for complex lipids and 2x10^6^ cells for fatty acids from each experimental group were aliquoted into 1.5mL Eppendorf tubes and washed twice with ice cold PBS. Supernatant was carefully removed, pellets were frozen with dry ice and stored at -80°C. Three biological samples were included for each condition. Further mass spectrometric analysis was performed at Weill Cornell Medicine Proteomics and Metabolomics core facility.

#### Complex lipid profiling

Lipids were extracted from samples using isopropanol according to a published method^66^.The extracts were clarified by centrifugation and dried down using a SpeedVac. The dried sample was reconstituted using acetonitrile/isopropanol/water 65:30:5 containing stable isotope labeled internal lipid standards (Splash Lipidomix, Avanti Polar Lipids) prior to LC-MS analysis. Chromatographic separation was performed on a Vanquish UHPLC system with a Cadenza CD-C18 3 µm packing column (Imtakt, 2.1 mm id x 150 mm) coupled to a Q Exactive Orbitrap mass spectrometer (Thermo Scientific) via an Ion Max ion source with a HESI II probe (Thermo Scientific). The mobile phase consisted of buffer A: 60% acetonitrile, 40% water, 10 mM ammonium formate with 0.1% formic acid and buffer B: 90% isopropanol, 10% acetonitrile, 10 mM ammonium formate with 0.1% formic acid. The LC gradient was as follows: 0–1.5 min, 32% buffer B; 1.5-4 min, 32-45% buffer B; 4-5 min, 45-52% buffer B; 5-8 min, 52-58% buffer B; 8-11 min, 58-66% buffer B; 11-14 min, 66-70% buffer B; 14-18 min, 70-75% buffer B; 21-25 min, isocratic 97% buffer B, 25-25.1 min 97-32% buffer B; followed by 5 min of re-equilibration of the column before the next run. The flow rate was 200 μL/min. A data-dependent mass spectrometric acquisition method was used for lipid identification. In this method, each MS survey scan was followed by up to 10 MS/MS scans performed on the most abundant ions. Data was acquired in positive and negative mode in separate runs. The following electrospray parameters were used: spray voltage 3.0 kV, heated capillary temperature 350 °C, HESI probe temperature 350 °C, sheath gas, 35 units; auxiliary gas 10 units. For MS scans: resolution, 70,000 (at m/z 200); automatic gain control target, 3e6; maximum injection time, 200 ms; scan range, 250-1800 m/z. For MS/MS scans: resolution, 17,500 (at 200 m/z); automatic gain control target, 1e5 ions; maximum injection time, 75 ms; isolation window, 1 m/z; NCE, stepped 20, 30 and 40. The LC-MS files were processed using MS-DIAL software (version 4.9) for lipid identification and relative quantitation. Peak intensities from complex lipid profiling were batch corrected.

#### Stable Isotope Tracing of Free Fatty Acids

Free fatty acids were extracted from cells using 90% methanol (LC/MS grade, Thermo Scientific). The extracts were clarified by centrifugation and dried down using a SpeedVac. The dried sample was reconstituted using 50% methanol prior to LC-MS analysis. Chromatographic separation was performed on a Vanquish UHPLC system (Thermo Scientific) with a Cadenza CD-C18 3 µm packing column (Imtakt, 2.1 mm id x 150 mm) coupled to a Q Exactive Orbitrap mass spectrometer (Thermo Scientific) via an Ion Max ion source with a HESI II probe (Thermo Scientific). The mobile phase consisted of buffer A: 5 mM ammonium acetate in water (LC/MS grade, Thermo Scientific) and buffer B: 5 mM ammonium acetate, 85% isopropanol (LC/MS grade, Thermo Scientific), 10% acetonitrile (LC/MS grade, Thermo Scientific), and 5% water. The LC gradient was as follows: 0–1.5 min, 50% buffer B; 1.5-3 min, 50-90% buffer B; 3-5.5 min, 90-95% buffer B; 5.5-10 min, 95% buffer B, followed by 5 min of re-equilibration of the column before the next run. The flow rate was 150 μL/min. MS data was acquired in negative mode. The following electrospray parameters were used: spray voltage 3.0 kV, heated capillary temperature 350 °C, HESI probe temperature 350 °C,sheath gas, 35 units; auxiliary gas 10 units. For MS scans: mass scan range, 140-1000 m/z; resolution, 70,000 (at m/z 200); automatic gain control target, 1e6; maximum injection time, 50 ms. MS data files were processed using El-MAVEN (v0.12.0). Identification of free fatty acids was based on accurate masses within 5 ppm and standard retention times. Relative quantitation was performed based on MS signal intensities.

#### Analysis

Data from both panels were processed through MetaboAnalyst 6.0 for statistical analysis with normalization by medium. Heatmaps were generated through the online platform. Normalized peak intensities were used to calculate log2 fold change, *p-* and *q-*values. Volcano plots were generated through R studio.

### MALDI TOF Imaging of Mouse Tissues

#### MALDI Sample Preparation

Mouse brains were harvested and snap-frozen for 5 min on an aluminum boat floating on liquid nitrogen as previously described^67^. The frozen tissue was cryosectioned at 10 μm thickness sections using CryoStar NX50 (Thermo Scientific, USA) at −15 °C set for both specimen head and the chamber. Sections of cerebrum around Bregma -2.3 mm and of cerebellum around -6.4 mm were collected according to Allen Mouse Brain Coronal Atlas plane 75 and 116 respectively. The tissue cryosections were gently transferred onto the pre-cooled conductive side of indium tin oxide (ITO)-coated glass slides (Bruker Daltonics,

Bremen, Germany) and thaw mounted using finger. Mounted cryosections on ITO slides were desiccated in vacuum for 45 min at room temperature, followed by matrix deposition using HXT M5 sprayer (HXT LLC., USA)^68,69^. Matrix NEDC (N-naphthylethylenediamine dihydrochloride) was used to detect fatty acids^70^, and a matrix solution of 10 mg/mL in isopropanol/water (70/30, v/v) was deposited at a flow rate of 0.05 ml/min and a nozzle temperature of 80 °C for 30 cycles with no drying between each cycle. A spray velocity of 1300 mm/min, track spacing of 2 mm, N2 gas pressure of 10 psi and flow rate of 3 L/min and nozzle height of 40 mm were used for matrixes.

#### MALDI Imaging Process

MALDI mass spectra were acquired in negative ion mode by MALDI time-of-light (TOF) mass spectrometer Autoflex (Bruker Daltonics, Germany). MS spectra were calibrated using red phosphorus as the standard for all experiments^68,69^. The laser spot diameters for two MALDI runs were focused to “Medium” modulated beam profile for both 40 μm and 130 μm raster width. The imaging data for each array position were summed up by 500 shots at a laser repetition rate of 500 Hz. The whole brain and lung tissue MALDI imaging at a resolution of 130 micrometers (µm). For the higher resolution scan at 40 µm, only the region of interest within each tissue section was imaged. The Spectra were acquired in the mass range from m/z 100 to 1000 with a low mass gate at 100 Da. Imaging data were recorded and processed using FlexImaging v3.0, and further analyzed using SCiLS (2015b). Ion images were generated with total ion count (TIC) normalization and a bin width of ±0.10 Da.

#### Metabolite Annotation

The spectra were interpreted manually, and analyte assignment was accomplished by comparing the data with previously published studies^70–72^. Initially, the possible element compositions for each ion were estimated using Metaboscape (Bruker Daltonics, Germany), based on its exact mass (with an expected mass accuracy of less than 5 ppm) and isotope patterns. These ions were then matched against several databases, including The Human Metabolome Database (HMDB), METLIN, LIPID MAPS, and a custom-made metabolite library (previous LC-MS data), to identify potential metabolites.

#### Segmentation Procedure for MALDI Data

Spatial segmentation of the MALDI-imaging data was carried out by clustering of spectra by their similarity with SCiLS segmentation tools. It can automatically generate a spatial segmentation map of the tissue section, where regions of similar metabolomic patterns were highlighted^73^. Initially, Find Peak tools was used to find those peaks inside a spectrum encoding relevant information, then a segmentation pipeline was executed on the acquired IMS data for unbiased hierarchical clustering^74^ with a minimal interval width of ±0.100 Da. The resulting clusters were examined for their correlation with histological features of the tissues manually. Subsequently, clusters representing relevant regions were selected for further analysis.

#### H&E Staining

MALDI slides were retrieved immediately after the experiment, washed with 95% ethanol for 30 sec and proceeded with standard H&E staining. The H&E images were acquired using Leica Aperio CS2 slide scanner and used as anatomical reference for mass spectra imaging.

## Acknowledgements

We are deeply grateful to the patients and families who inspire and support our daily work. We would like to thank Weill Cornell Medicine Proteomics and Metabolomics Core Facility, MSKCC molecular cytology and flow cytometry core facilities for their support in this research. We would also like to thank Areej Niaz and Dr. Rinat Abzalimov for technical support on MALDI MSI.

This research is supported by NCI 1 R01 CA245499-01A1, the Geoffrey Beene Foundation, Alan and Sandra Gerry Metastasis and Tumor Ecosystems Center, Pershing Square Sohn Cancer Research Alliance, all to A.B. The STARR Cancer Consortium and the Pew Charitable Trusts, to A.B. and N.K. NIH R01DK136013588, NIH R01NS12654 and PSC-CUNY Research Award Program support the work of Y.H., and L.X.

## Author contributions

X.T. and A.B. conceptualized and designed the study, wrote the manuscript. X.T. conducted the experiments, analyzed and compiled the data. J.R., M.J.L., J.S. A.R.N. H.W., A.O., and A.Y.L.W. assisted with mouse experiments. S.P.O. and B.M. assisted with *in vitro* experiments. J.B. and B.P. performed and analyzed the mouse CSF targeted metabolomics under N.K. supervision. L.X. performed and analyzed the mouse tissue MALDI mass spectrum imaging under Y.H. supervision. K.C., R.E. and I.O.G. carried out with clinical sample annotation and human CSF processing. M.S. generated and validated the LLC LeptoM *Nr1h2/3* DKO and *Rxra/b/g* TKO cell lines under T.Z. supervision. P.J.H. and R.K. analyzed the bulk ATAC sequencing datasets. R.M. provided and the human patient autopsy samples. A.B. provided oversight of all experimental design and analysis.

## Declaration of interests

J.R. and A.B. are inventors on the following provisional US patent applications: 63/449,817, and 63/449,823, and international patent application: PCT/US24/18343. A.B. holds an unpaid position on the Scientific Advisory Board for Evren Scientific and is an inventor on the following patents: 62/258,044, 10/413,522, and 63/052,139.

---

**Supplementary Figure 1:**
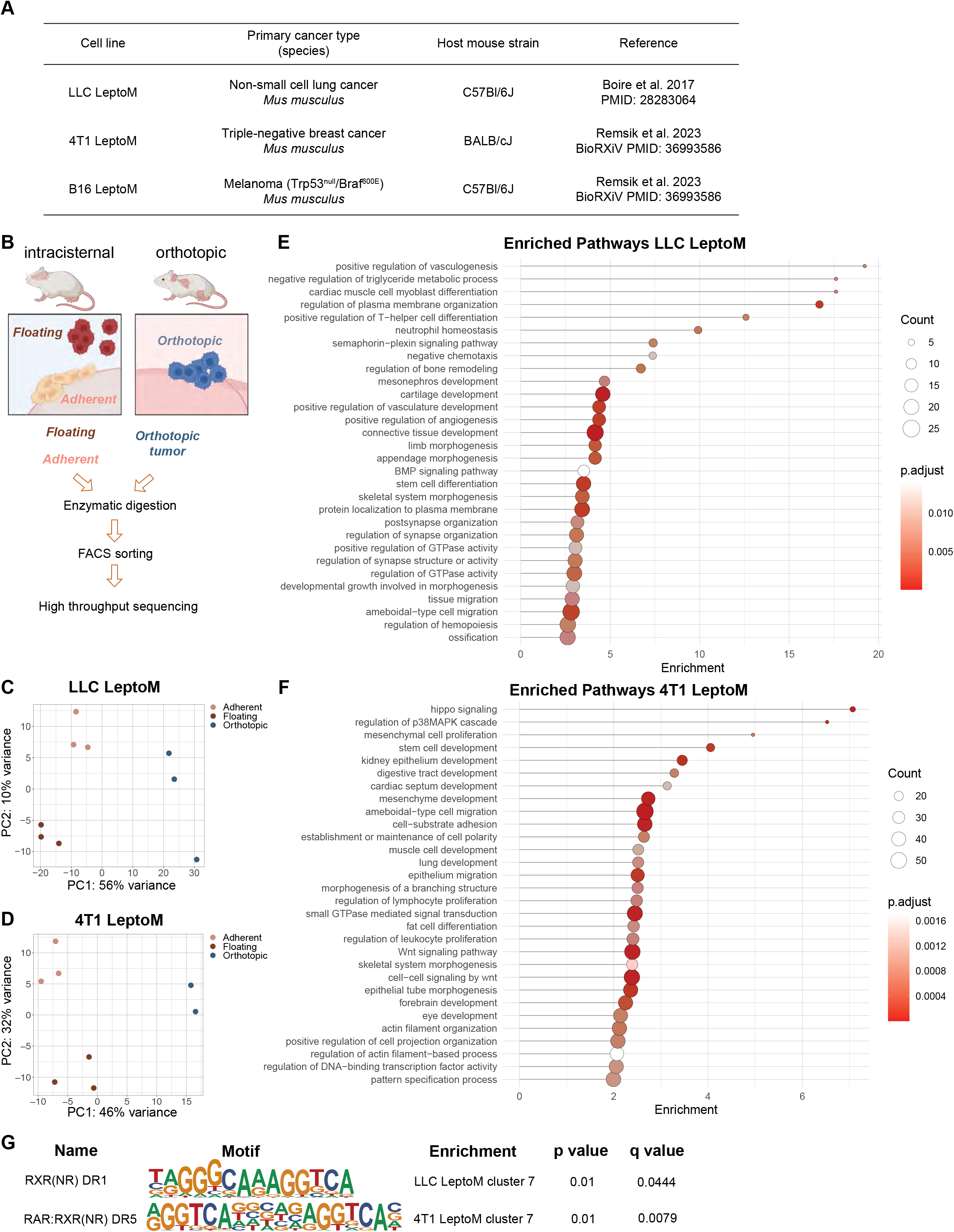
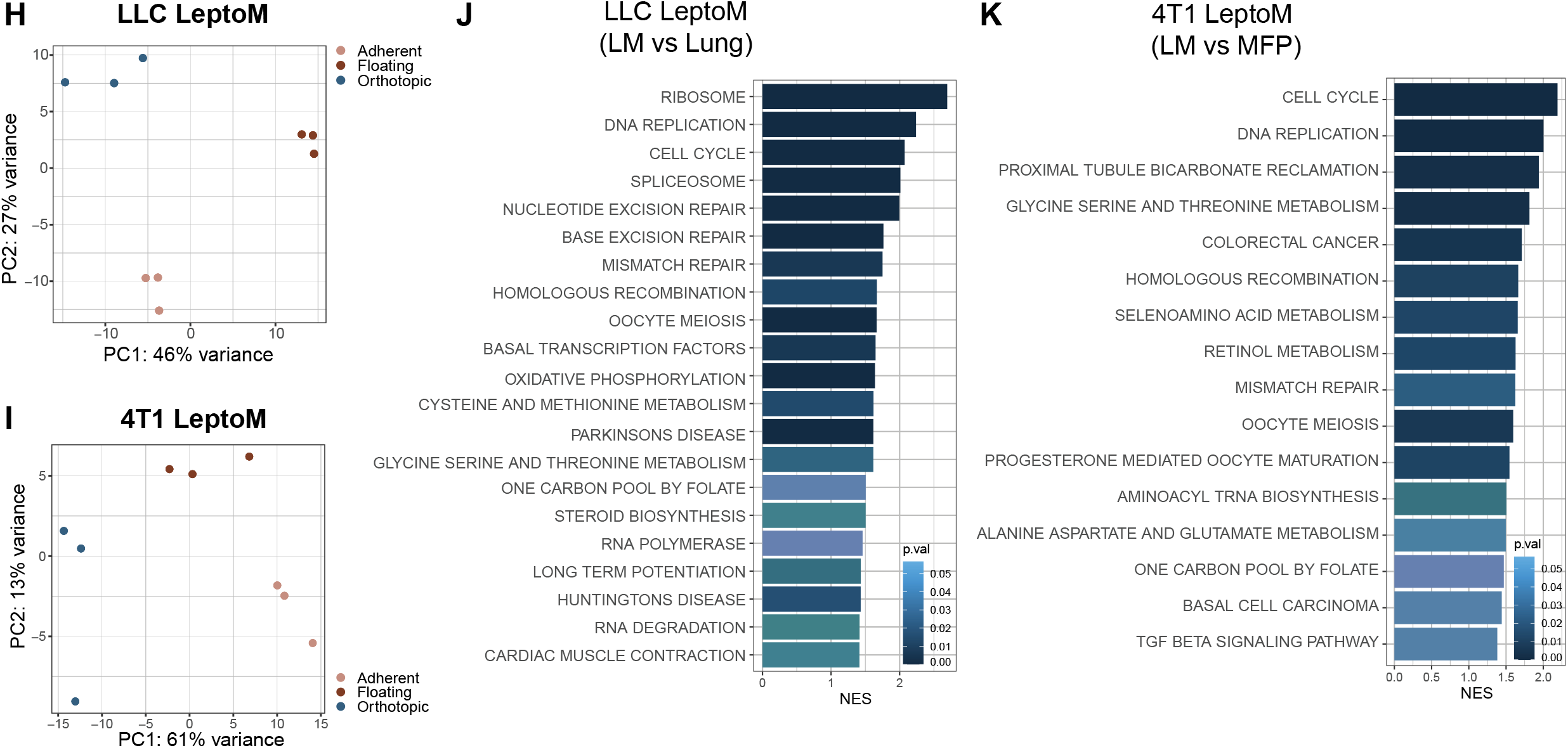
Freshly sorted LLC and 4T1 LeptoM from distinct microenvironments revealed unique chromatin and transcriptomic landscapes. (A) Overview of leptomeningeal metastatic cell lines (LeptoM) used in this paper and their background information. (B) Schematic drawing of the bulk ATAC- and RNA-sequencing experimental design for LLC and 4T1 LeptoM cells. Detailed procedure can be found in Methods. (C-D) Principal Component Analysis (PCA) using ATAC-sequencing signals for LLC (C) and 4T1 (D) LeptoM based on peak normalization method. LLC LeptoM contain three replicates for each group. 4T1 LeptoM contain three replicates for adherent and floating fractions, and two replicates for orthotopic group. (E-F) Differentially accessible genes comparing leptomeningeal-residing cells to orthotopic sites were selected for pathway analysis. Enrichment in LLC (E) and 4T1 (F) LeptoM are depicted with a highlight of developmental pathways. (G) RXR-related motifs are enriched in the open regions from leptomeningeal-residing cells in LLC and 4T1 LeptoM cluster 7 (please refer to Figure 1B-C) based on ATAC-seq signals. (H-I) PCA of bulk RNA sequencing of LLC (H) and 4T1 (I) LeptoM. LLC LeptoM and 4T1 LeptoM both contain three replicates for each group. Plots are generated with DESeq package. (J-K) Gene Set Enrichment Analysis (GSEA) performed comparing leptomeningeal- residing cells to orthotopic cells in LLC (J) and 4T1 (K) LeptoM. KEGG pathways are extracted and plotted.

**Supplementary Figure 2:**
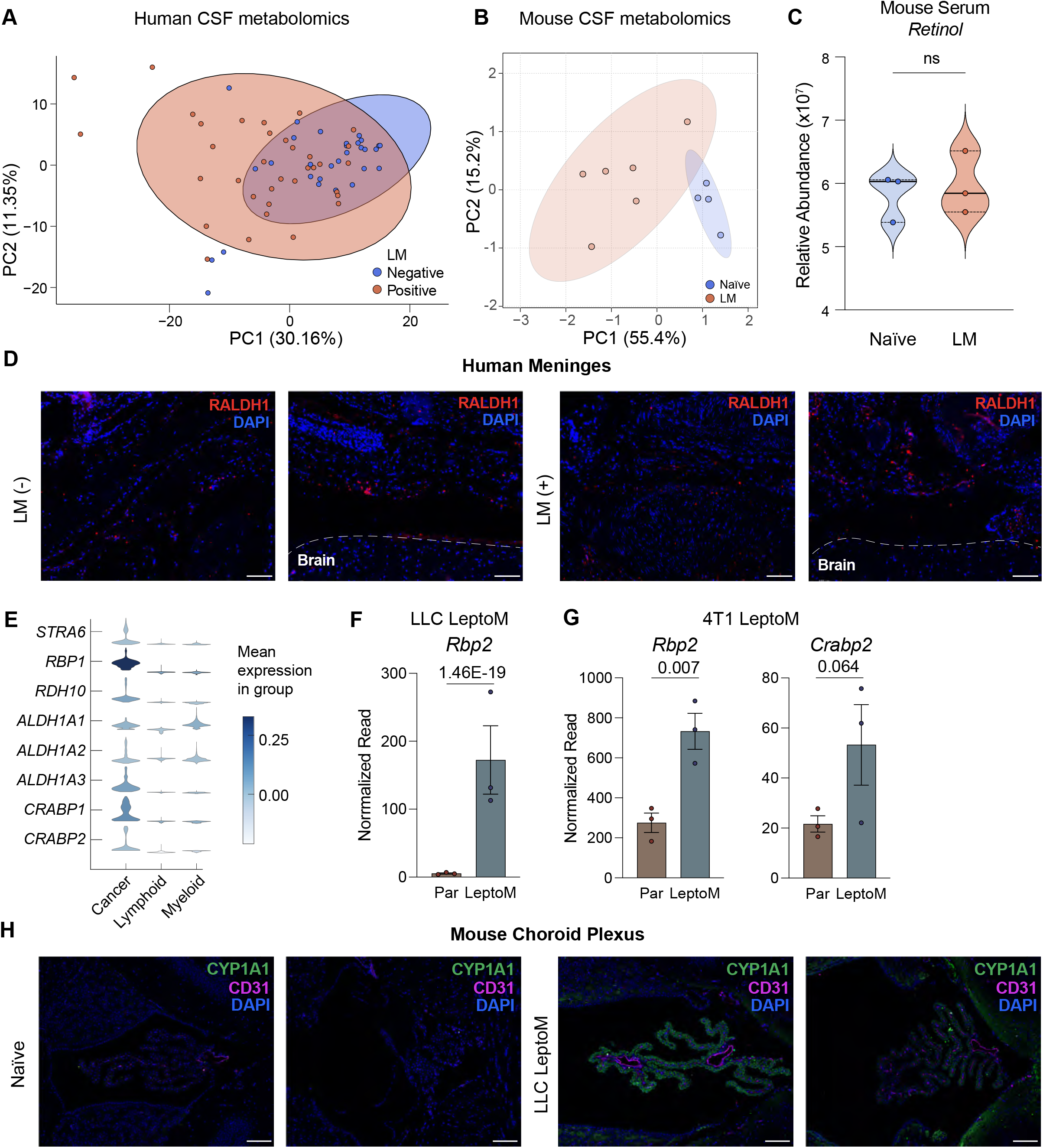
Cancer cells in the leptomeninges demonstrate evidence of upregulated retinol metabolism. (A) PCA plot of untargeted metabolomics from human patient CSF without LM diagnosis (n=30) or with new LM diagnosis (n=35). Total of 554 compounds of known identity are depicted. (B) PCA plot of targeted metabolomics from mouse CSF from naïve mice or mice injected with 2,000 LLC LeptoM intracisternally. 15 polar metabolites were detected and used for normalization of *all-trans* retinol in the same samples. (C) *All-trans* retinol level was detected on LC-MS/MS in the serum from naïve mice (n=3) or mice injected with 2,000 LLC LeptoM intracisternally (n=3). Retinol standard was used for identification of retinol peak. Peak areas were normalized to a panel of polar metabolites (please refer to Figure S2B and Methods for details). Mouse serum was collected on day 14 after injection. *p*-value represents the *F*-test to compare population variance. (D) Immunofluorescent staining of RALDH1 and DAPI on meninge sections using human autopsy samples. Sections from two patients without LM and two sections from one patient with LM diagnosis were selected. White dash lines mark the edge of brain. Scale bar = 100μm. (E) Stacked violin plot generated from previously published scRNA-seq on 10X platform of five human LM samples^6^. Genes encoding retinol uptake, mobilization and metabolism are expressed in the LM cancer cell population. (F) Normalized reads of *retinol binding protein 2 (Rbp2)* extracted from previously published *in vitro* bulk RNA-seq comparing LLC Par and LeptoM^11^. Error bars represent SEM. Statistical tests are *q*-values. (G) Normalized read of *Retinol binding protein 2 (Rbp2) and Cellular retinoic acid binding protein 2 (Crabp2)* extracted from previously published *in vitro* bulk RNA-seq comparing 4T1 Par and LeptoM^12^. Error bars represent SEM. Statistical tests are *q*-values. (H) Immunofluorescent staining of CYP1A1, CD31(marker for endothelial cells) and DAPI in 3^rd^ ventricle choroid plexi on mice brain sections collected from naïve mouse (n=2) or mouse intracisternally injected with 2,000 LLC LeptoM-mCherry cells (n=2). Scale bar = 100μm.

**Supplementary Figure 3:**
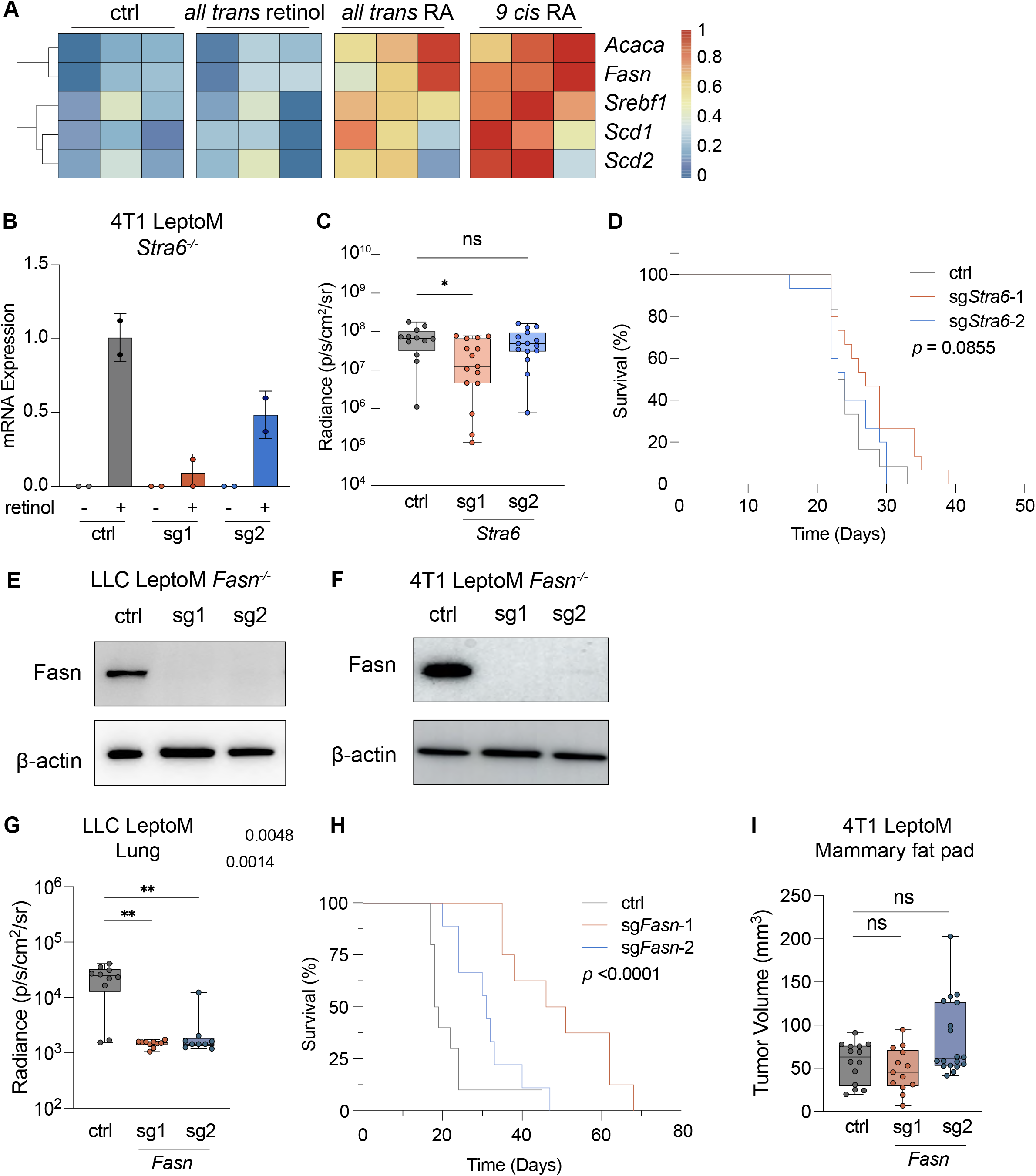
RA-activated *de novo* FA synthesis supports cancer cell growth in the leptomeningeal space. (A) RT-qPCR assaying *de novo* FA biosynthesis genes (*Srebf1, Fasn, Acaca, Scd1, Scd2)* using Taqman probes in B16 LeptoM cells upon stimulation with 1μM *all-trans* retinol, 1μM *all-trans* RA, 1μM *9-cis* RA, or vehicle control (DMSO). Housekeeping genes Gapdh and *β*-actin were used as endogenous control. Stimulation in lipid-depleted media were normalized to B16 LeptoM DMSO-treated control in complete media. Plot contains three independent experiments each with duplicates. (B) RT-qPCR confirming *Stra6* gene expression loss in 4T1 LeptoM Stra6 knockout cells with and without 1μM retinol stimulation *in vitro*. Housekeeping gene *Gapdh* was used as endogenous control. Error bar = SD. (C) 4T1 LeptoM transduced with vector control or two independent small guide RNAs targeting *Stra6* were injected intracisternally into recipient mice; tumor growth was monitored by BLI weekly. Day 14 BLI was plotted as average radiance on log10 scale. Three biological experiments were performed. Ctrl: n=12; sg*Stra6*-1: n=15; sg*Stra6-*2: n=15. Error bars represent SEM. * represents *p*=0.0328, *p*-value calculated by non- parametric one-way ANOVA (Kruskal-Wallis test). (D) Kaplan-Meier survival curve of mice from (C). Log-rank test was used for *p*-value. (E) Western blot confirmation for loss of Fasn in LLC LeptoM cell lines (sg1, sg2). Two biological experiments were performed independently, one is shown. *β*-actin was used as loading control. (F) Western blot confirmation for loss of Fasn in 4T1 LeptoM cell lines (sg1, sg2). Two biological experiments were performed independently, one is shown. *β*-actin was used as loading control. (G) 20,000 of LLC LeptoM vector control, sg*Fasn*-1 or sg*Fasn*-2 cells were injected into lung parenchyma of the recipient mice, tumor growth was monitored by BLI weekly. Day 14 BLI was plotted as average radiance. Two biological experiments were performed. Ctrl: n=10; sg*Fasn*-1: n=10; sg*Fasn*2: n=9. Error bars represent SEM. ** represents *p*<0.005, calculated by non-parametric one-way ANOVA (Kruskal-Wallis test). (H) Kaplan-Meier survival curve of mice from (G). Log-rank test was used for *p*-value. (I) 50,000 of 4T1 LeptoM vector control, sg*Fasn*-1 or sg*Fasn*-2 cells were injected into mammary fat pad of the recipient mice, tumor growth was measured by Caliper twice every week. Day 14 tumor volume was calculated. Two biological experiments were performed. Ctrl: n=14; sg*Fasn*-1: n=13; sg*Fasn*2: n=18. Error bars represent SEM, *p*- value calculated by non-parametric one-way ANOVA (Kruskal-Wallis test).

**Supplementary Figure 4:**
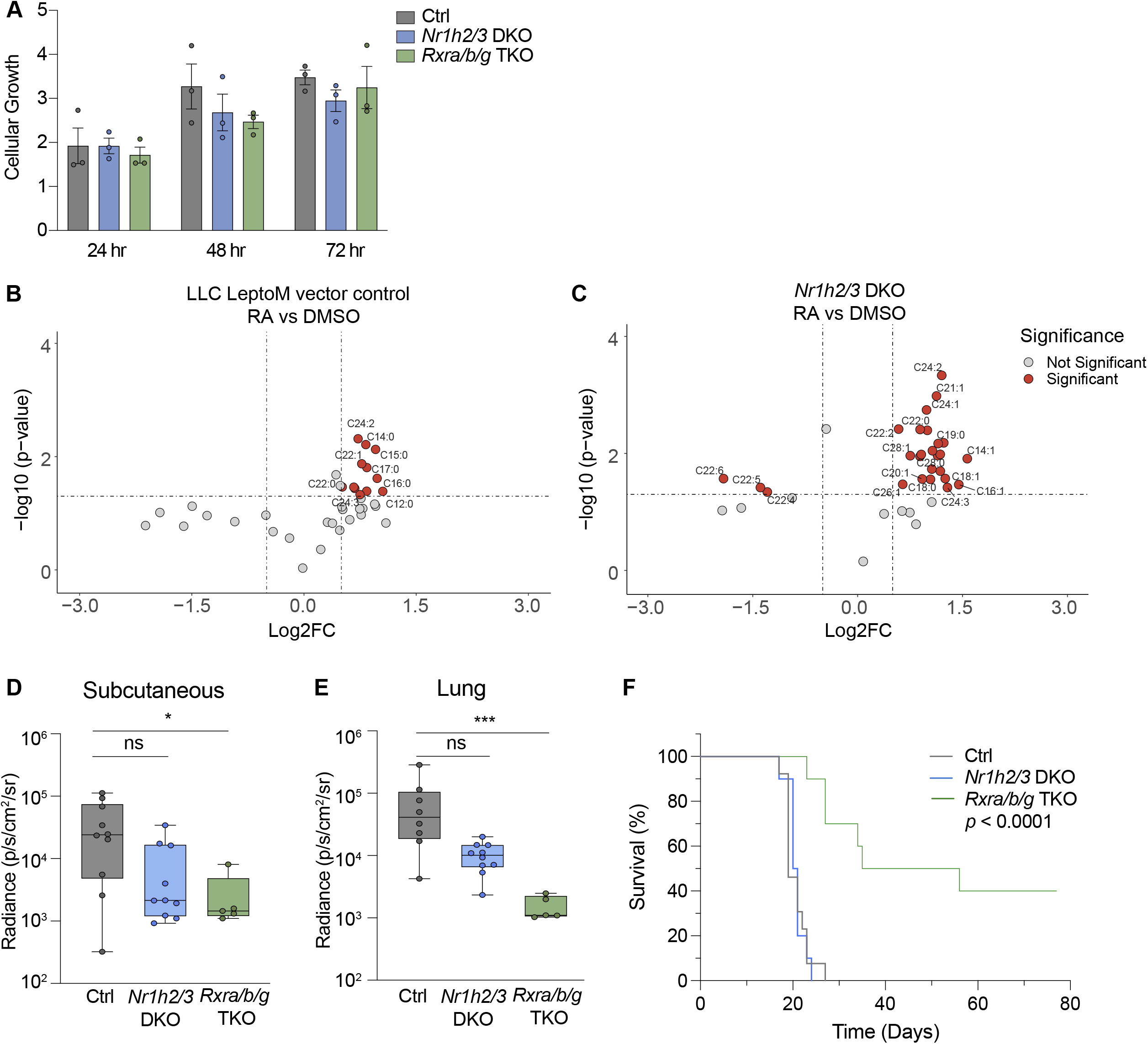
Characterizing LLC LeptoM *Nr1h2/3* DKO and *Rxra/b/g* TKO. (A) Cellular proliferation of LLC LeptoM vector control, *Nr1h2/3* DKO or *Rxra/b/g* TKO were assessed by colorimetric Cell Titer 96 Aqueous assay 24, 48 and 72hrs after seeding *in vitro*. Readings were normalized to each cell line 0hr seeding control. Three independent biological experiments were performed with four technical replicates each. (B) Volcano plot of LC-MS/MS based intracellular free FA lipidomic analysis of LLC LeptoM vector control treated 1μM *9-cis* RA compared to DMSO. Significantly changed free FA were annotated and colored in red. Significant cutoff: |log2 fold change| > 0.5, *p*- value < 0.05. (C) Volcano plot of LC-MS/MS based intracellular free FA lipidomic analysis of LLC LeptoM *Nr1h2/3* DKO treated 1μM *9-cis* RA compared to DMSO. Significantly changed free FA were annotated and colored in red. Significant cutoff: |log2 fold change| > 0.5, *p*- value < 0.05. (D) 50,000 of LLC LeptoM vector control, *Nr1h2/3* DKO or *Rxra/b/g* TKO cells were injected into subcutaneous fat of the recipient mouse flanks, tumor growth was monitored by BLI weekly. Day 14 BLI was plotted as average radiance normalized to day 0. Two biological experiments were performed. Ctrl: n=10; *Nr1h2/3* DKO: n=10, *Rxra/b/g* TKO: n=5. Error bars represent SEM, *p*-value calculated by non-parametric T-test (Mann- Whitney test), * represents *p*=0.0301. (E) 20,000 of LLC LeptoM vector control, *Nr1h2/3* DKO or *Rxra/b/g* TKO cells were injected into lung parenchyma of the recipient mice, tumor growth was monitored by BLI weekly. Day 14 BLI was plotted as average radiance normalized to day 0. Two biological experiments were performed. Ctrl: n=8; *Nr1h2/3* DKO: n=10, *Rxra/b/g* TKO: n=5. Error bars represent SEM, *p*-value calculated by non-parametric T-test (Mann-Whitney test), *** represents *p*=0.0002. (F) Kaplan-Meier survival curve of mice from (E). Ctrl: n=13; *Nr1h2/3* DKO: n=10, *Rxra/b/g* TKO: n=10. *p*-value calculated by log-rank test between LLC LeptoM vector control and *Rxra/b/g* TKO.

**Supplementary Figure 5:**
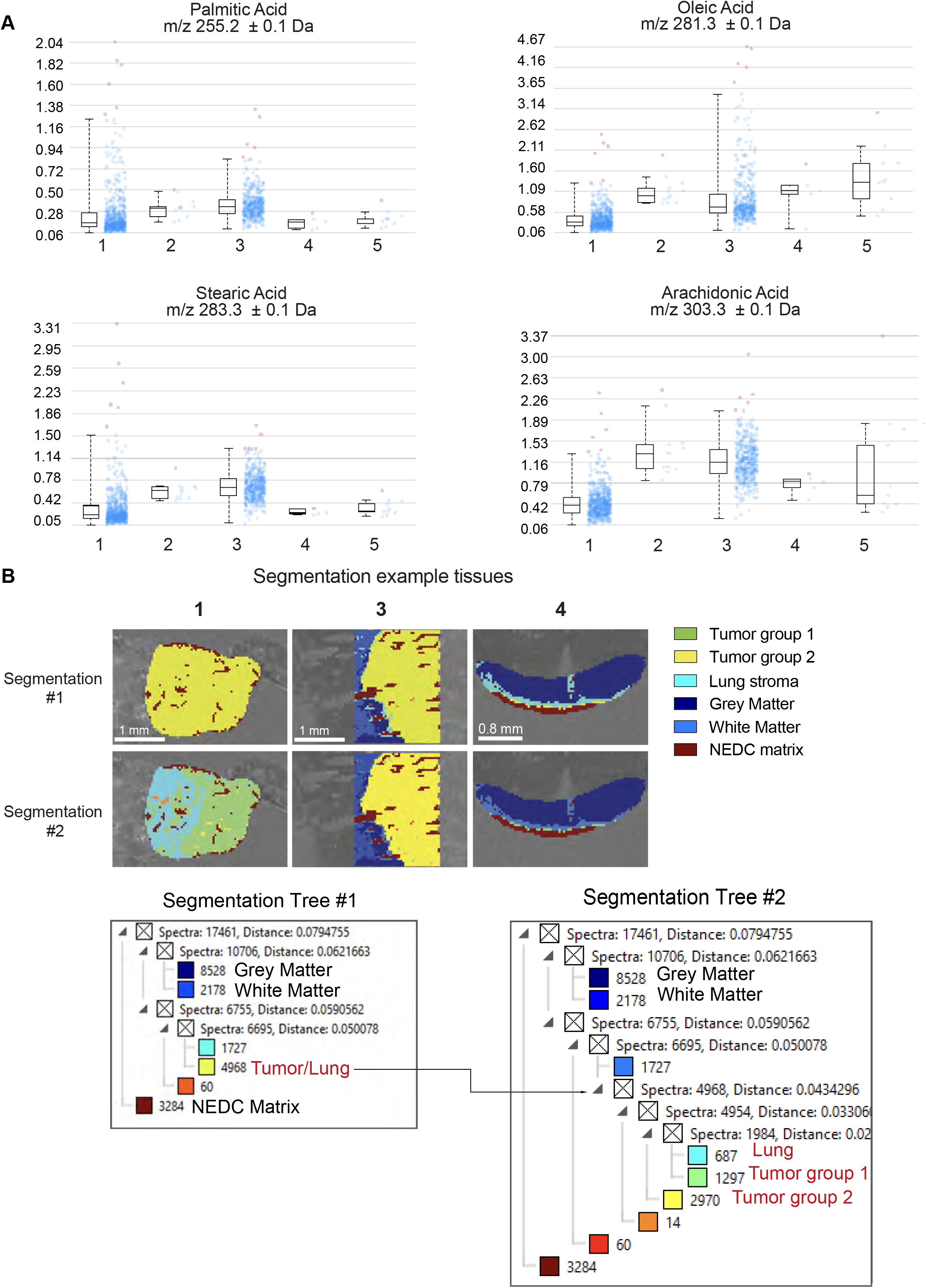
Spatial metabolomics reveals distinct metabolic spectrum of LeptoM cells *in vivo*. (A) Intensity box plots of tumor region fatty acids plotted in Figure 5B generated by SCiLS software after 130μm segmentation. Each line in the box represents mean peak intensity, upper and lower box boundary represent third and second quartiles. Each point represents the normalized peak intensity in the pixel identified as cancer cells after segmentation. (B) Representative image of LLC LeptoM injected lung (tissue 1), brain (tissue 3) and LLC LeptoM *Nr1h2/3* DKO brain (tissue 4) at 40μm scan to show segmentation #1 and segmentation #2. Both segmentation trees are presented, arrow shows at that level segmentation is carried further. Tissues identified based on H&E staining and Allen Brain Atlas are annotated on the segmentation tree.

**Supplementary Figure 6:**
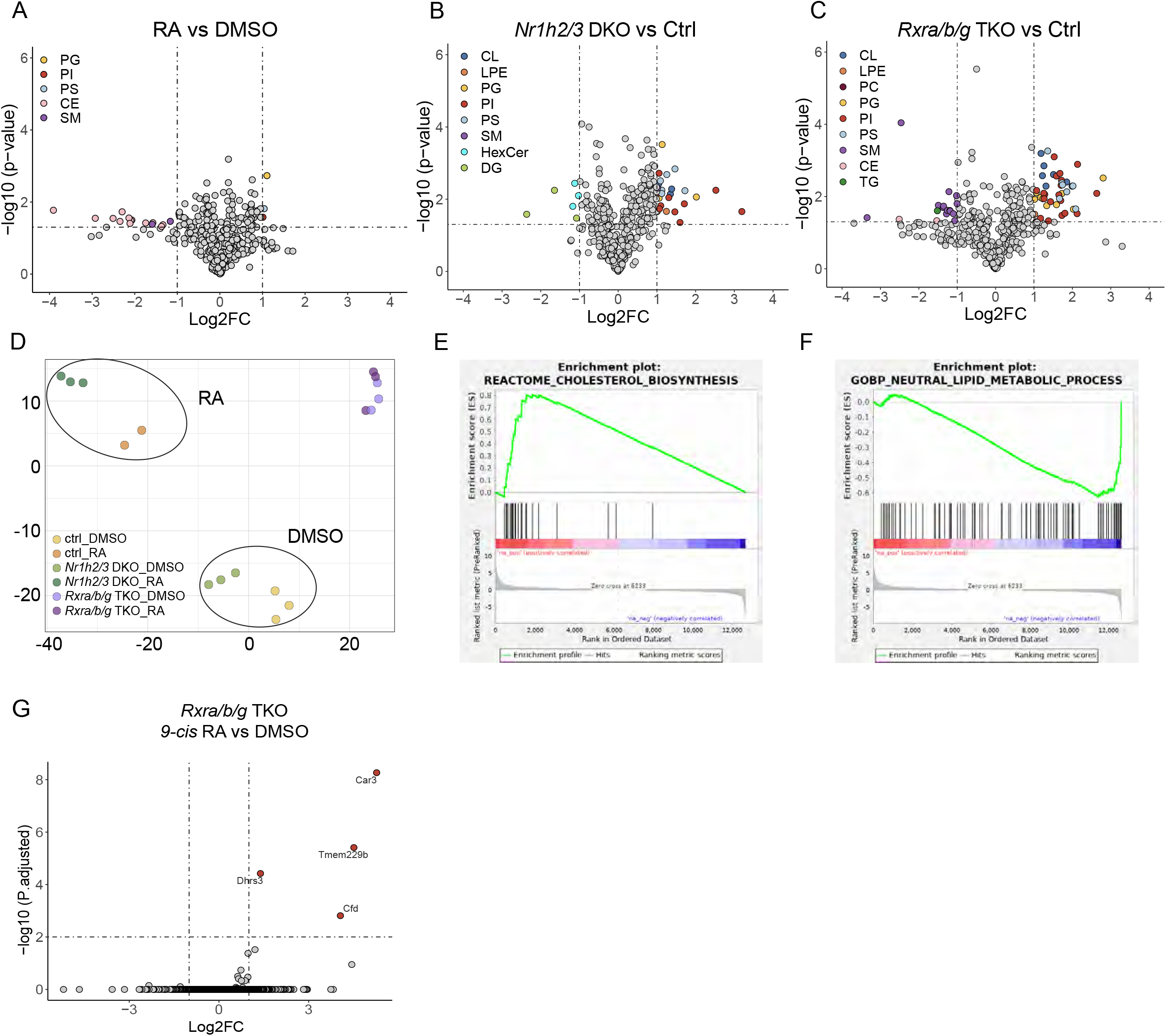
Transcriptome and metabolome analysis of RXR- mediated signaling in LeptoM cells. (A) Volcano plot of LC-MS/MS based intracellular complex lipid profiling of LLC LeptoM vector control treated 1μM *9-cis* RA compared to DMSO. Significantly changed lipids were colored by category. Significant cutoff: |log2 fold change| > 1, *p*-value < 0.05. PG: phosphatidylglycerol; PI: phosphatidylinositol; PS: phosphatidylserine; CE: Cholesteryl ester; SM: sphingomyelin. (B) Volcano plot of LC-MS/MS complex lipid profiling comparing LLC LeptoM *Nr1h2/3* DKO to LLC LeptoM vector control both treated with DMSO. Significantly changed complex lipid species were colored by category. Significant cutoff: |log2 fold change| > 1, *p*-value < 0.05. CL: cardiolipins; LPE: Lysophosphatidylethanolamine; HexCer: hexosylceramide; DG: diacylglycerol. (C) Volcano plot of LC-MS/MS complex lipid profiling comparing LLC LeptoM *Rxra/b/g* TKO to LLC LeptoM vector control both treated with DMSO. Significantly changed complex lipid species were colored by category. Significant cutoff: |log2 fold change| > 1, *p*-value < 0.05. PC: phosphatidylcholine; TG: Triglyceride. (D) PCA for bulk RNA-seq with LLC LeptoM vector control, *Nr1h2/3* DKO and *Rxra/b/g* TKO cells treated with DMSO vehicle control or 1μM *9-cis* RA *in vitro*. (E) GSEA analysis of upregulated genes in LLC LeptoM vector control treated 1μM *9-cis* RA versus DMSO. Reactome pathway cholesterol biosynthesis is shown in the plot with NES = 1.87, FDR = 0.046. (F) GSEA analysis of downregulated genes in LLC LeptoM vector control treated 1μM *9- cis* RA versus DMSO. Gene Ontology Biological Processes (GOBP) pathway neutral lipid metabolic process is shown in the plot with NES = -2.08, FDR = 0.011. (G) Volcano plot of bulk RNA-seq comparing LLC LeptoM *Rxra/b/g* TKO treated with 1μM *9-cis* RA to DMSO. Significant cutoff: |log2 FC| > 1, *P-adjusted* < 0.01.

| Patient Identifier | Gender | Age | CSF protein (mg/dL) | CSF glucose (mg/dL) | Primary diagnosis | LM | CTC |
| --- | --- | --- | --- | --- | --- | --- | --- |
| 018-1 | F | 72 | 92 | 88 | Breast Cancer | Negative | 1 |
| 025-1 | F | 59 | 41 | 88 | Breast Cancer | Positive | NA |
| 029-2 | F | 52 | 64 | 29 | Breast Cancer | Positive | >200 |
| 057-1 | F | 50 | 208 | 8 | Lung Cancer | Positive | >200 |
| 059-1 | F | 48 | 123 | 43 | Breast Cancer | Positive | 65 |
| 061-1 | F | 61 | 34 | 27 | Lung Cancer | Positive | >200 |
| 062-1 | M | 72 | 42 | 51 | Lung Cancer | Negative | 0 |
| 071-1 | F | 72 | 67 | 41 | Lung Cancer | Positive | >200 |
| 072-1 | F | 49 | 38 | 67 | Lung Cancer | Positive | 44 |
| 074-1 | F | 53 | 66 | 23 | Breast Cancer | Positive | >200 |
| 085-1 | F | 56 | 53 | 52 | Lung Cancer | Positive | >200 |
| 100-1 | F | 38 | 160 | 52 | Breast Cancer | Positive | 46 |
| 127-1 | M | 52 | NA | NA | Lung Cancer | Positive | NA |
| 137-1 | F | 61 | 54 | 40 | Lung Cancer | Positive | 32 |
| 145-2 | M | 57 | 50 | 58 | Lung Cancer | Positive | 134 |
| 151-1 | F | 57 | 32 | 60 | Lung Cancer | Negative | 0 |
| 160-1 | F | 79 | 51 | 73 | Breast Cancer | Negative | 0 |
| 164-1 | M | 68 | 119 | 50 | Lung Cancer | Positive | 189 |
| 166-1 | F | 77 | 55 | 59 | Breast Cancer | Positive | 14 |
| 167-2 | F | 55 | 656 | <5 | Breast Cancer | Positive | >200 |
| 169-1 | F | 38 | 679 | 70 | Breast Cancer | Positive | 7 |
| 170-1 | F | 34 | 41 | 54 | Breast Cancer | Positive | >200 |
| 171-1 | F | 55 | 61 | 58 | Breast Cancer | Negative | 0 |
| 172-1 | F | 59 | 26 | 52 | Lung Cancer | Negative | 0 |
| 173-1 | M | 50 | NA | NA | Lung Cancer | Negative | NA |
| 174-1 | F | 50 | 56 | 81 | Lung Cancer | Positive | >200 |
| 181-1 | M | 51 | 104 | 71 | Melanoma | Positive | NA |
| 182-1 | F | 57 | 74 | 11 | Breast Cancer | Positive | >200 |
| 190-2 | F | 40 | 88 | 44 | Breast Cancer | Positive | 183 |
| 186-1 | F | 54 | 47 | 55 | Breast Cancer | Negative | 0 |
| 266-1 | F | 67 | 211 | 12 | Lung Cancer | Positive | >200 |
| 198-1 | F | 62 | 330 | 68 | Breast Cancer | Positive | 33 |
| 202-1 | F | 55 | 58 | 55 | Breast Cancer | Negative | 1 |
| 205-1 | M | 75 | 92 | 104 | Melanoma | Negative | 0 |
| 206-1 | M | 65 | 46 | 60 | Lung Cancer | Negative | 0 |
| 207-1 | F | 61 | 28 | 52 | Melanoma | Positive | NA |
| 222-1 | F | 76 | 33 | 55 | Lung Cancer | Positive | >200 |
| 282-1 | F | 68 | 91 | 59 | Breast Cancer | Negative | 1 |
| 246-1 | M | 41 | 26 | 68 | Melanoma | Negative | NA |
| 302-1 | F | 74 | 43 | 80 | Lung Cancer | Negative | NA |
| 304-1 | F | 73 | 37 | 56 | Lung Cancer | Negative | 0 |
| 351-1 | M | 66 | 36 | 59 | Lung Cancer | Negative | 1 |
| 360-1 | F | 46 | NA | NA | Breast Cancer | Negative | NA |
| 364-1 | F | 46 | 26 | 68 | Breast Cancer | Negative | 0 |
| 384-1 | F | 82 | 1078 | 42 | Melanoma | Positive | NA |
| 390-1 | F | 40 | 179 | 77 | Melanoma | Negative | 0 |
| 438-1 | M | 63 | 35 | 57 | Melanoma | Negative | 0 |
| 460-1 | F | 69 | 582 | 44 | Lung Cancer | Positive | 98 |
| 472-1 | F | 48 | 31 | 58 | Lung Cancer | Negative | 0 |
| 474-1 | M | 71 | 50 | 58 | Breast Cancer | Negative | NA |
| 484-1 | M | 61 | 50 | 58 | Melanoma | Positive | NA |
| 485-1 | M | 60 | 104 | 49 | Melanoma | Positive | NA |
| 486-1 | M | 82 | 234 | 56 | Melanoma | Positive | NA |
| 487-1 | F | 53 | 74 | 111 | Melanoma | Positive | NA |
| 488-1 | M | 77 | 150 | 57 | Melanoma | Positive | 0 |
| 489-1 | M | 92 | 239 | 44 | Melanoma | Negative | NA |
| 490-1 | M | 21 | 105 | 61 | Melanoma | Positive | 0 |
| 491-1 | M | 60 | 57 | 77 | Melanoma | Negative | NA |
| 492-1 | M | 78 | 111 | 100 | Melanoma | Negative | 0 |
| 493-1 | F | 47 | 23 | 61 | Melanoma | Negative | NA |
| 494-1 | M | 75 | 58 | 87 | Melanoma | Negative | 0 |
| 495-1 | M | 75 | 56 | 59 | Melanoma | Negative | NA |
| 496-1 | M | 24 | 26 | 66 | Melanoma | Negative | NA |
| 497-1 | F | 43 | 31 | 78 | Breast Cancer | Positive | 173 |
| 283-1 | F | 47 | 151 | 69 | Lung Cancer | Negative | 0 |

| Gender | Age | Primary diagnosis | LM |
| --- | --- | --- | --- |
| M | 61 | Prostate cancer | Negative |
| F | 65 | (ER+/HER2+) Breast Cancer | Negative |
| F | 4 | Stage IV Neuroblastoma | Negative |
| M | 27 | Desmoplastic Small Round Cell Tumor (DSRCT) | Negative |
| F | 45 | (ER/PR/HER2+) stage IV breast cancer | Positive |
| M | 49 | (P53+, EGFR-) Non-Small Cell Lung Cancer (NSCLC) | Positive |
| F | 55 | Metastatic breast cancer | Positive |

